# Geometric model incorporating prey’s turn and predator attack endpoint explains multiple preferred escape trajectories

**DOI:** 10.1101/2020.04.27.049833

**Authors:** Yuuki Kawabata, Hideyuki Akada, Ken-ichiro Shimatani, Gregory N. Nishihara, Hibiki Kimura, Nishiumi Nozomi, Paolo Domenici

## Abstract

The escape trajectory (ET) of prey – measured as the angle relative to the predator’s approach path – plays a major role in avoiding predation. Previous geometric models predict a single ET; however, many species show highly variable ETs with multiple preferred directions. Although such a high ET variability may confer unpredictability to avoid predation, the reasons why animals prefer specific multiple ETs remain unclear. Here, we constructed a novel geometric model that incorporates the time required for prey to turn and the predator’s position at the end of its attack. The optimal ET was determined by maximizing the time difference of arrival at the edge of the safety zone between the prey and predator. By fitting the model to the experimental data of fish *Pagrus major* we show that the model can clearly explain the observed multiple preferred ETs. By using the same model, we were able to explain different patterns of ETs empirically observed in other species (e.g., insects and frogs): a single preferred ET and multiple preferred ETs at small (20–50°) and large (150–180°) angles from the predator. Our results open new avenues of investigation for understanding how animals choose their ETs from behavioral and neurosensory perspectives.

## Introduction

When exposed to sudden threatening stimuli such as ambush predators, most prey species initiate escape responses that include turning swiftly and accelerating away from the threat. The escape responses of many invertebrate and lower vertebrate species are controlled by giant neurons that ensure a short response time [1]. Many previous studies have focused on two behavioral traits that are fundamental for avoiding predation: when to escape (i.e., flight initiation distance, which is measured as the distance from the predator at the onset of escape) and where to escape [i.e., escape trajectory (ET), which is measured as the angle of escape direction relative to the stimulus direction] [2]. Previous studies have investigated the behavioral and environmental contexts affecting these variables [3–8], because they largely determine the success or failure of predator evasion [9–13], and hence the fitness of the prey species. A large number of models on how animals determine their flight initiation distances have been formulated and tested by experiments [2]. Although a number of models have also been developed to predict animal ETs [4, 14, 15], there are still some unanswered questions about how the variability of the observed ETs is generated.

Previous geometric models predict a single ET that depends on the relative speeds of the predator and the prey [4, 14–16], and additionally, predator’s turning radii and sensory-motor delay in situations where the predator can adjust its strike path [17–19]. However, these models do not explain the complex ET distributions reported in empirical studies on various taxa of invertebrates and lower vertebrates (reviewed in [20]). Whereas some animals exhibit unimodal ET patterns that satisfy the geometric models (e.g., [21]), many animals show multimodal ETs within a limited angular sector (esp., 90–180°) (e.g., [4, 5, 22]). To explore the discrepancy between the predictions of the models and empirical data, some researchers have hypothesized mechanical/sensory constraints [20, 23] and unpredictability, in line with the idea of a protean response that does not allow predators to adopt counter-strategies [23–27]. Although these hypotheses, together with the previous geometric models, can explain the ET variability within a limited angular sector, the reasons why animals prefer specific multiple ETs remain unclear.

In previous geometric models, the prey was assumed to instantaneously escape in any direction, irrespective of the prey’s initial body orientation relative to the predator’s approach path (hereafter, initial orientation) [4, 14, 15]. However, additional time is required for changing the heading direction (i.e., turn); therefore, a realistic model needs to take into account that the predator can approach the prey while the prey is turning [12]. Additionally, in previous models, attacking predators were assumed to move for an infinite distance at a constant speed [4, 14, 15]. However, the attacks of many real predators, especially ambush ones, end at a certain distance from initial positions of the prey [28–30]. Therefore, we constructed a geometric model that incorporates two additional factors: the time required for the prey to turn and the endpoint of the predator attack. First, using a fish species as a model, we tested whether our model could predict empirically observed multimodal ETs. Second, by extending the model, we tested whether other patterns of empirical ETs could be predicted: unimodal ETs and multimodal ETs directed at small (20–50°) and large (150–180°) angles from the predator’s approach direction. The biological implications resulting from the model and experimental data are then discussed within the framework of predator-prey interactions.

## Model

We revised previous models proposed by Domenici [15, 31] and by Corcoran & Conner [18] (Fig. 1A, B). Other previous models [4, 14, 16, 19] made predictions similar to those of Domenici’s model or those of Corcoran’s model, although they used different theoretical approaches. In Domenici’s model, the predator with a certain width (i.e., the width of a killer whale’s tail used as a weapon to catch prey) directly approaches the prey, and the prey (the whole body) should enter the safety zone before the predator reaches that entry point. In this model, the prey can instantaneously escape in any direction, and the predation threat moves linearly and infinitely. Corcoran’s model is based on the same principle as Domenici’s model, but includes the concept that the predator (i.e., a bat) can adjust the approach path up to its minimum turning radius. Thus, Domenici’s model can be regarded as a special case of Corcoran’s model when the turning radius of the predator is infinitely large. These models are based on the escape response of the horizontal plane, which is realistic for many fish species as well as terrestrial and benthic species that move on substrates. They can also be applied to aerial animals such as moths escaping from bats because many predator-prey interactions are approximately two-dimensional in a local spatial scale [18, 32].

**Fig. 1.**
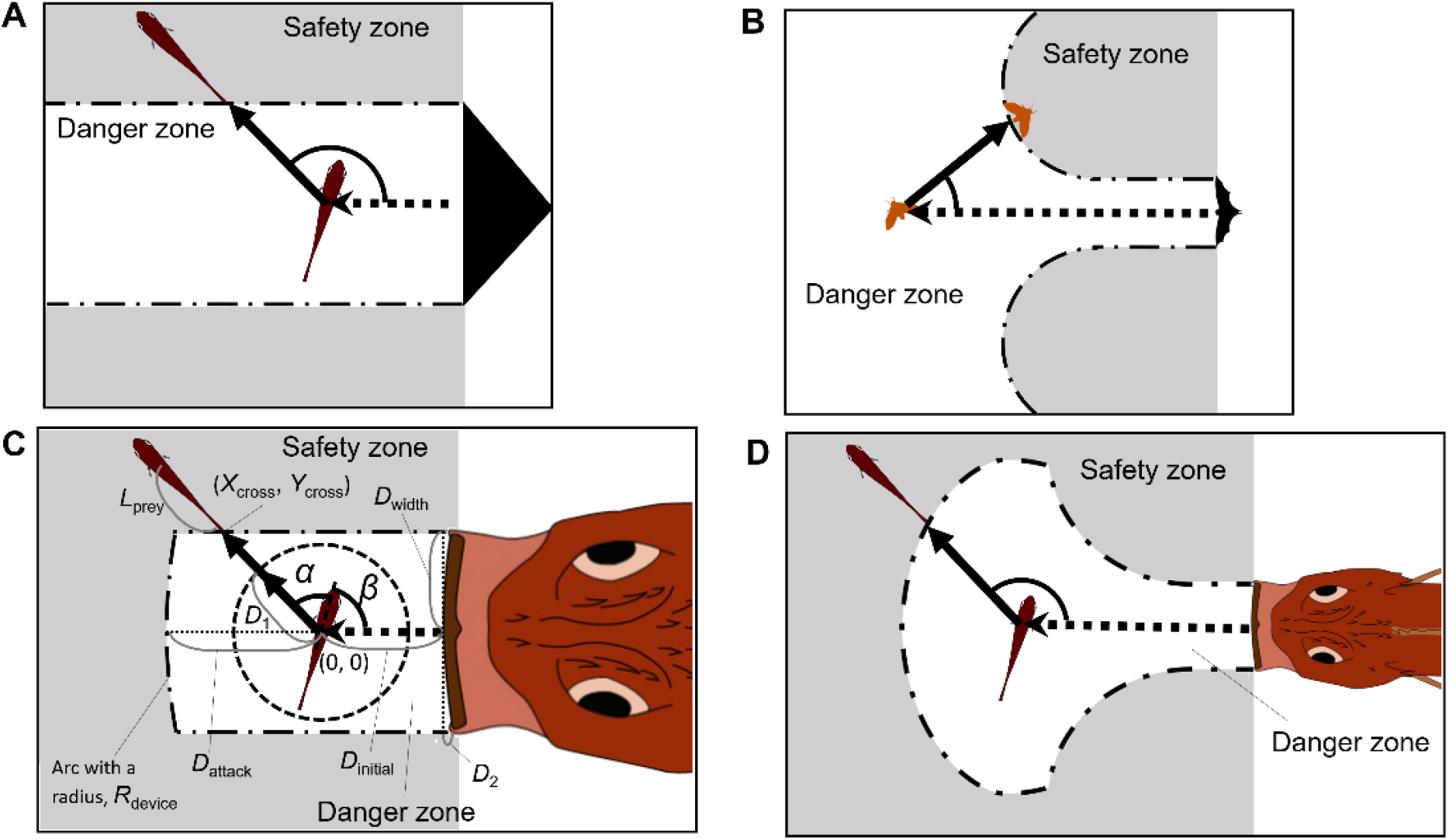
Proposed geometric models for animal escape trajectories. (A) A previous geometric model proposed by Domenici [15]. The predation threat with a certain width (the tail of a killer whale, represented by the black triangle) directly approaches the prey, and the prey should reach the safety zone (a grey area) outside the danger zone (white area) before the threat reaches that point. In this model, the prey can instantaneously escape in any direction, and the predation threat moves linearly and infinitely. (B) A previous geometric model proposed by Corcoran & Conner [18]. This model is based on the same principle as Domenici’s model, but includes the concept that the predator (i.e., a bat) can adjust the approach path up to its minimum turning radius. (C) Two factors are added to Domenici’s model: the endpoint of the predator attack, and the time required for the prey to turn. See Table 1 and the text for details of the definitions of the variables and mathematical formulas. (D) Two factors are added to Corcoran’s model: the endpoint of the predator attack, and the time required for the prey to turn. See SI Appendix for details of the definitions of the variables and mathematical formulas. Note that Domenici’s model (A) and its modified model (C) can be regarded as special cases of Corcoran’s model (B) and its modified model (D), respectively, when the turning radius of the predator is infinitely large.

In our new models (Fig. 1C, D), two factors are added to the previous Domenici’s and Corcoran’s models: the time required for the prey to turn and the endpoint of the predator attack. We assume that a prey with a certain initial orientation *β* (spanning 0–180°, where 0° and 180° correspond to being attacked from front and behind, respectively) evades a sudden predation threat. Most prey species respond to the attack by turning at an angle *α* and the ET results from the angular sum of *α* and *β*. ETs from the left and right sides were pooled and treated as though they were stimulated from the right side (Fig. S1; See “Definition of Angles” in SI Appendix for details). Hereafter, in the main text, we explain the modification of Domenici’s model (a special case of Corcoran’s model) because the data on previously published predator-prey experiments on the same species of prey and predator in our experiment [12] show that the predator does not adjust the strike path during the attack [Fig. S2, adjusted angle=1.0±6.6° (mean±s.d.), n=5], and thus the number of parameters to estimate can be reduced. See SI Appendix and Fig. S3 for details of the modified version of Corcoran’s model.

**Table 1.** Methods for determining parameter values

| Symbol | Description | Value | Method |
| --- | --- | --- | --- |
| $D_{\text{width}}$ | the half-width of the predator capture device (e.g., mouth) | 18 mm | measured directly from the dummy predator (a sacrificed individual) |
| $R_{\text{device}}$ | the radius of the predator's capture device at the moment of attack, which is approximated as an arc | 199 mm | measured directly from the dummy predator (a sacrificed individual) |
| $D_1$ | the displacement from the initial position of prey where it was assumed to end the fast-start phase | 15 mm | estimated from the escape kinematics of prey in the experiment |
| $U_{\text{prey}}$ | the prey speed after the displacement of $D_1$, which is assumed to be constant | 1.04 m s<sup>-1</sup> | estimated from the escape kinematics of prey in the experiment |
| $T_1(\vert \alpha\vert )$ | the time required for a displacement of $D_1$ from the initial position of the prey, given by a function of turn angle $\vert \alpha\vert $ | Fig. 3A | estimated from the escape kinematics of prey in the experiment |
| $D_{\text{attack}}$ | the distance between the prey's initial position and the tip of the predator capture device at the end of the predator attack | 35 mm | optimized by comparing the model outputs with experimental data |
| $U_{\text{pred}}$ | the predator speed, which is assumed to be constant | 1.54 m s<sup>-1</sup> | optimized by comparing the model outputs with experimental data |

When the prey’s center of mass (CoM) at the onset of its escape is located at point (0, 0), the trajectory of the CoM (*X*_prey_, *Y*_prey_) is given by:

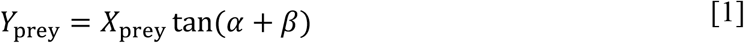

The edge of the safety zone is determined by the half-width of the predator capture device (e.g., mouth) *D*_width_, the distance between the prey’s initial position and the tip of the predator capture device at the end of the predator attack *D*_attack_, and the shape of the predator’s capture device at the moment of attack, which is approximated as an arc with a certain radius, *R*_device_. The projection of the predator’s capture device edge along the edge of the sideways safety zone *D*_2_ can be expressed as:

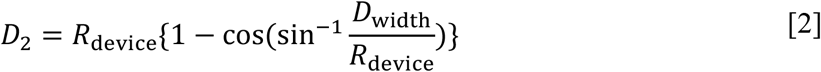

The ET toward the upper-left corner of the danger zone *θ*_corner_ can be expressed as:

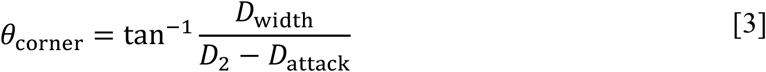

The x and y coordinates of the safety zone edge (*X*_safe_, *Y*_safe_) are given by:

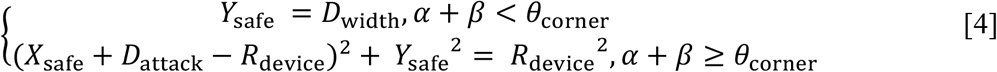

From equations [1] to [4], the x and y coordinates of the crossing point of the escape path and the safety zone edge (*X*_cross_, *Y*_cross_) are given by a function of *D*_width_, *D*_attack_, *R*_device_, and *α*+*β*.

The prey can escape from the predator when the time required for the prey to enter the safety zone (*T*_prey_) is shorter than the time required for the predator’s capture device to reach that entry point (*T*_pred_). Therefore, the prey is assumed to maximize the difference between the *T*_pred_ and *T*_prey_ (*T*_diff_). To incorporate the time required for the prey to turn, *T*_prey_ was divided into two phases: the fast-start phase, which includes the time for turning and acceleration (*T*_1_), and the constant speed phase (*T*_2_). This assumption is consistent with the previous studies [33–35] and was supported by our experiment (See Fig. S5). Therefore:

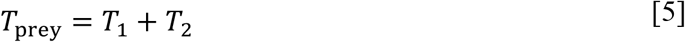

For simplicity, the fish was assumed to end the fast-start phase at a certain displacement from the initial position in any *α* (*D*_1_; the radius of the dotted circle in Fig. 1C) and to move at a constant speed *U*_prey_ to cover the rest of the distance (toward the edge of the safety zone 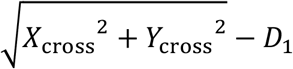, plus the length of the body that is posterior to the center of mass *L*_prey_). Because a larger |*α*| requires further turning prior to forward locomotion, which takes time [33, 36], and the initial velocity after turning was dependent on |*α*| in our experiment (See Fig. 3B), *T*_1_ is given by a function of |*α*| [*T*_1_(|*α*|)]. Therefore, *T*_prey_ can be expressed as:

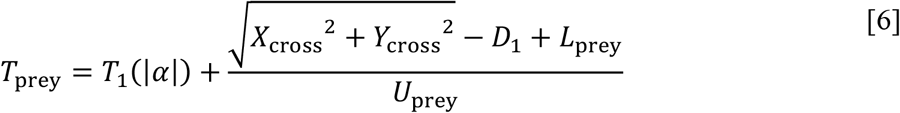

**Fig. 2.**
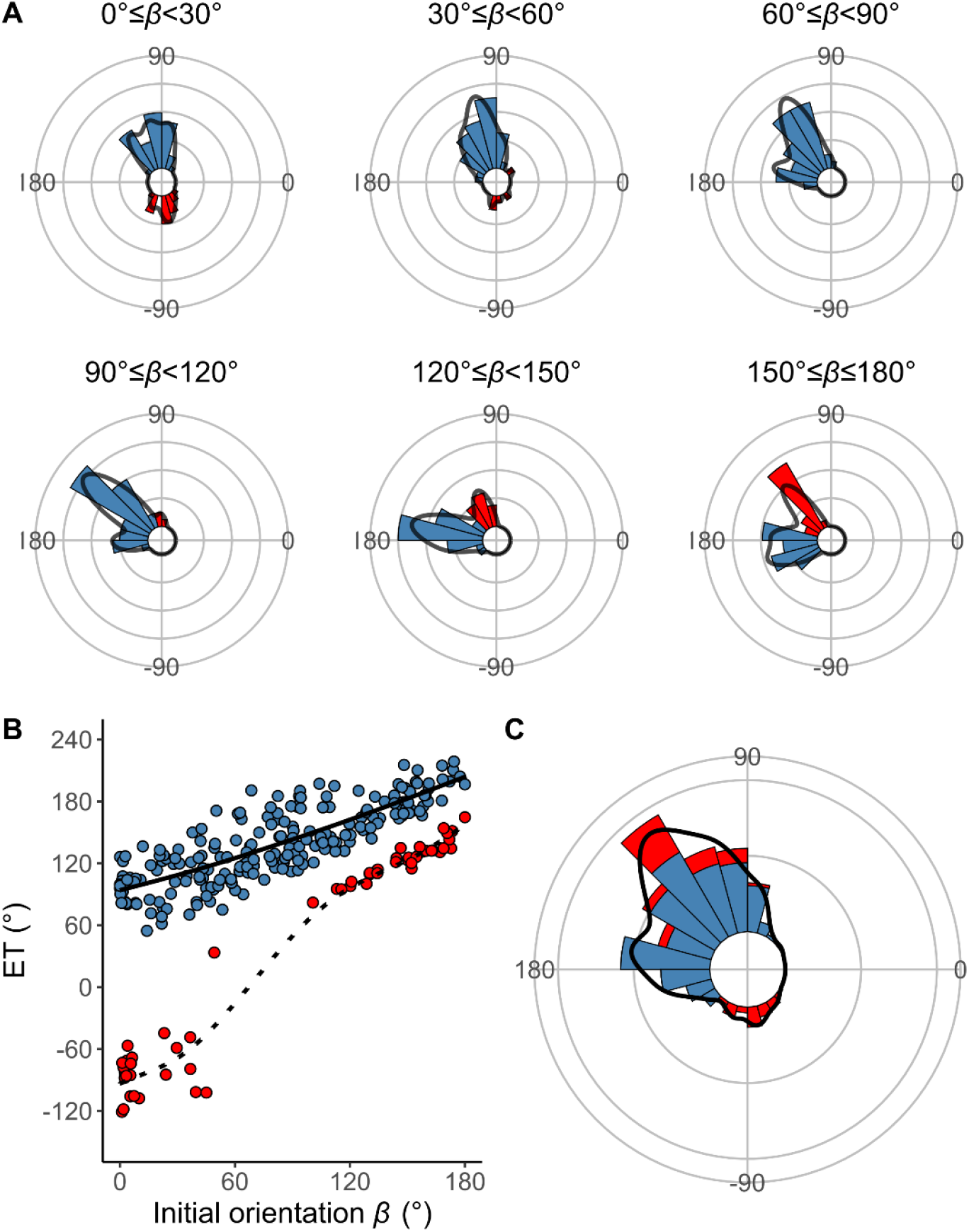
Results of the experiments of *Pagrus major* attacked by a dummy predator (i.e., a cast of *Sebastiscus marmoratus*). (A) Circular histograms of escape trajectories (ETs) in 30° initial orientation *β* bins. Solid lines are estimated by the kernel probability density function. Concentric circles represent 5% of the total sample sizes, the bin intervals are 15°, and the bandwidths of the kernel are 50. (B) Relationship between initial orientation and ET. Different colors represent the away (blue) and toward (red) responses. Solid and dotted lines are estimated by the generalized additive mixed model (GAMM). (C) Circular histogram of ETs pooling all the data shown in A. Solid lines are estimated by the kernel probability density function. Concentric circles represent 5% of the total sample sizes, the bin intervals are 15°, and the bandwidths of the kernel are 50. The predator’s approach direction is represented by 0° [n=264 (208 away and 56 toward responses) from 23 individuals].

**Fig. 3.**
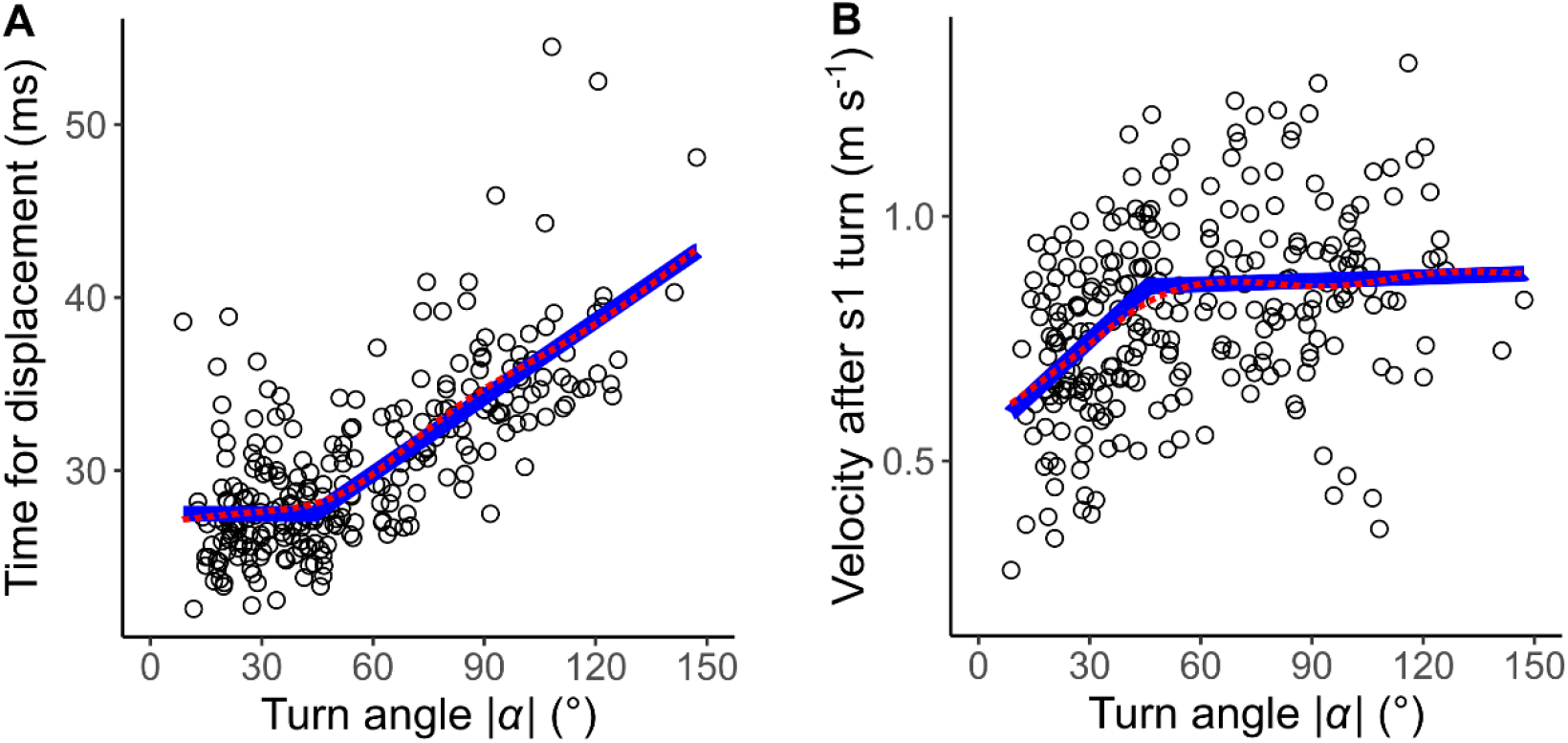
The relationship between the absolute value of the turn angle |*α*| and time-distance variables. (A) Relationship between |*α*| and the time required for a displacement of 15 mm from the initial position of the prey. (B) Relationship between |*α*| and the initial velocity after stage 1 turn. Solid blue lines are estimated by the piecewise linear regression model, and red dotted lines are estimated by the generalized additive mixed model (GAMM) (n=264 from 23 individuals).

*T*_pred_ can be expressed as:

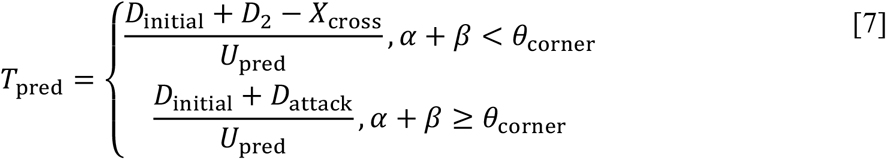

where *D*_initial_ is the distance between the prey and the predator at the onset of the prey’s escape response (i.e., the flight initiation distance or reaction distance), and *U*_pred_ is the predator speed, which is assumed to be constant. From equations [5] to [7], *T*_diff_ can be calculated as:

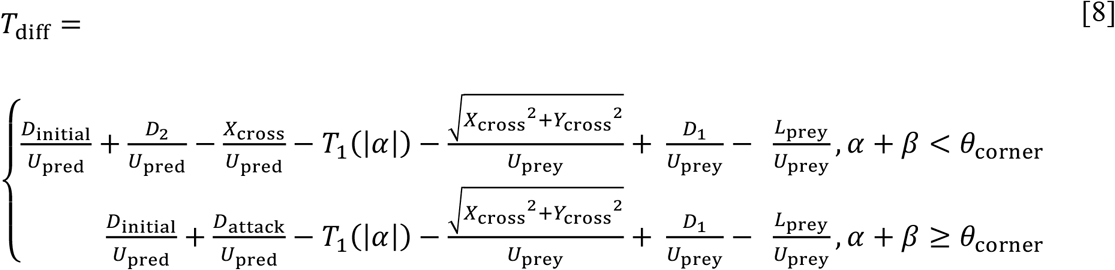

Because 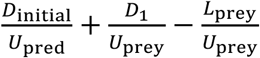 are independent of *α* and *β* we can calculate the relative values of 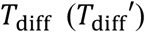 in response to the changes of *α* and *β* from:

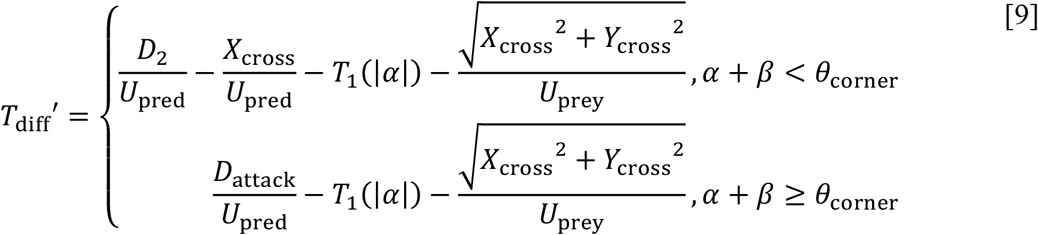

Because *X*_cross_ and *Y*_cross_ are dependent on *D*_width_, *D*_attack_, and *R*_device_ as well as *α* + *β* and *D*_2_ is dependent on *D*_width_ and *R*_device_, we can calculate 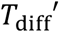 in response to the changes of *α* and *β* from *D*_1_, *D*_width_, *D*_attack_, *R*_device_, *U*_prey_, *U*_pred_, and *T*_1_(|*α*|) Given that the escape success is assumed to be dependent on 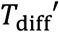, the theoretically optimal ET can be expressed as:

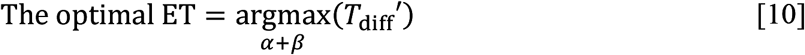

## Results

### Experimental Results

*P. major* exhibited a typical C-start escape response (Fig. S1), which consists of the initial bend (stage 1), followed by the return tail flip (stage 2), and continuous swimming or coasting (stage 3) [37, 38]. Figure 2 shows the effect of the initial orientation *β* on the ETs. As was done in previous studies [20, 39, 40], the away (contralateral) and toward (ipsilateral) responses, defined as the first detectable movement of the fish oriented either away from or toward the predator, were analyzed separately. When the initial orientation was small (i.e., the prey was attacked head-on; Fig. 2A; 0°≤*β*<30°), two peaks in the ET distribution were observed: a larger peak at around 100° (away response) and a smaller one at around −80° (toward response). As the initial orientation increases (Fig. 2A; 30°≤*β*<60°), the peak at around −80° disappeared. As the initial orientation further increases beyond 60°, another peak appeared at around 170° (Fig. 2A). When the initial orientation was large (i.e., the prey was attacked from behind; Fig. 2A; 150°≤*β*≤180°), there were two similar-sized peaks in the ET at around 130° (toward response), and 180–200° (away response). There were significant effects of initial orientation on the ET in both the away and the toward responses [away: generalized additive mixed model (GAMM), *F*=214.81, *P*<0.01, n=208; toward: GAMM, *F*=373.92, *P*<0.01, n=56]. There were significant effects of initial orientation on the turn angle *α* in away and toward responses (Fig. S4; away: GAMM, *F*=90.88, *P*<0.01, n=208; toward: GAMM, *F*=42.48, *P*<0.01, n=56). In the overall frequency distribution of ETs pooling the data on all initial orientations and both toward and away responses, there were two large peaks at 120–130° and 170–180°, and one small peak at around −80° (Fig. 2C). These 3 peaks were confirmed by the Gaussian mixture model analysis [41], where we fitted 1–9 Gaussian curves to the ETs, and selected the most parsimonious model based on the Akaike Information Criterion (AIC) (Table S1).

There were no significant effects of predator speed on the ET and |*α*| in either the toward or the away responses (ET, away: GAMM, *F*=0.01, *P*=0.93, n=208; ET, toward: GAMM, *F*=0.05, *P*=0.82, n=56; |*α*|, away: GAMM, *F*=0.01, *P*=0.93, n=208; |*α*|, toward: GAMM, *F*=0.05, *P*=0.82, n=56). There were no significant effects of predator speed [slow (from the minimum to the 33.3% quantile): 0.13~0.93 m/s; and fast (from the 66.7% quantile to the maximum): 1.29~1.88 m/s] on the variations of ETs and |*α*| in all 30° initial orientation bins (Levene’s test, *W*=0.01~3.57, *P*=0.07~0.91, n=13~32).

### Determination of Parameter Values

To predict the relationship between the ET (*α*+*β*) and the relative time difference *T*_diff_ in each initial orientation (*β*) by the geometric model, we needed *D*_width_, *R*_device_, *D*_1_, *U*_prey_, *T*_1_(|*α*|), *D*_attack_, and *U*_pred_. The methods for determining parameter values are summarized in Table 1. *D*_width_ and *R*_device_ were determined from the mouth shape of the predator (the sacrificed specimen for making the dummy predator) when fully opened, which were 18 and 199 mm, respectively. *D*_1_, *U*_prey_, and *T*_1_(|*α*|) were directly estimated by analyzing the escape responses of the prey. Because we have no previous knowledge about the values of *U*_pred_ and *D*_attack_ that the prey regards as dangerous, optimal values of *U*_pred_ and *D*_attack_ were determined iteratively by comparing model outputs with observed ETs. These optimal values were checked afterward with the data from previously published predator-prey experiments on the same species of prey and predator [12].

The distance of the fast-start phase (*D*_1_) was regarded as 15 mm based on the relationship between displacement and velocity of the prey in the experiments (Fig. S5), where the velocity increased up to about 15 mm of displacement from the initial position, beyond which it plateaus; over the 15 mm displacement from the initial position, there were no significant differences in the mean velocity between any combinations of 3-mm intervals in any 30° |*α*| bins (Fig. S5; paired *t*-test with Bonferroni’s correction, all *P*=1.00, n=23). There were significant effects of |*α*| on the time for a displacement of 15 mm from the initial position (GAMM, *F*=70.31, *P*<0.01, n=264) and on the mean velocity during the displacement (GAMM, *F*=69.49, *P*<0.01, n=264). However, there were no significant effects of |*α*| on the time required for a displacement of 15 to 30 mm from the initial position (GAMM, *F*=1.52, *P*=0.22, n=264) and on the mean velocity during the displacement (GAMM, *F*=0.89, *P*=0.27, n=264). Therefore, the time required for the prey to turn was incorporated into the model by analyzing the relationship between |*α*| and the time required for a displacement of 15 mm. The mean velocity of the prey during the constant phase *U*_prey_ was estimated to be 1.04 m s^−1^, based on the experimental data. Because the cut-off distance might affect the overall results of the study, we have repeated all the statistical analyses (See Tables 2, 3, and the text below for results with a cut-off distance of 15 mm) with cut-off distances of 10 and 20 mm and confirmed that the overall results are insensitive to the changes (Tables S2 and S3).

**Table 2.**
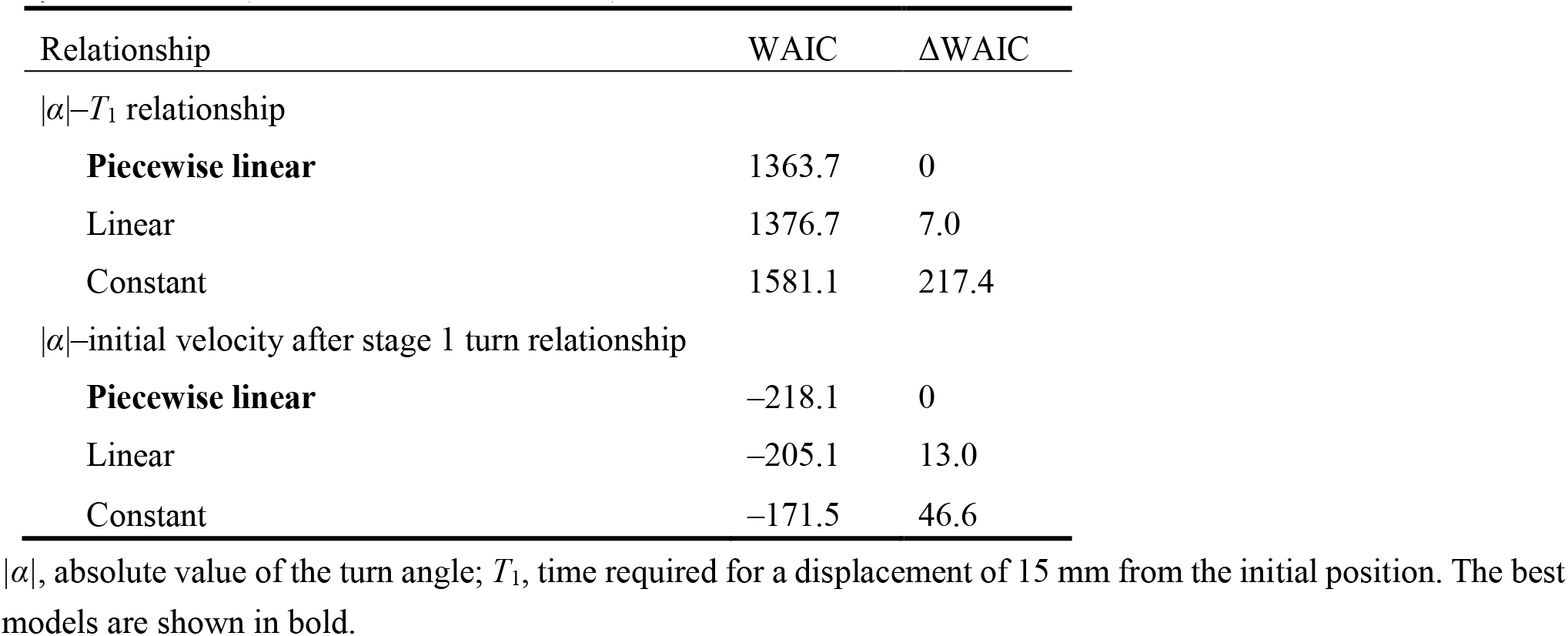
Widely applicable or Watanabe–Akaike information criterion (WAIC) for each model in the hierarchical Bayesian models (n=264 from 23 individuals)

| Relationship | WAIC | ΔWAIC |
| --- | --- | --- |
| $\vert \alpha\vert $–$T_1$ relationship | | |
| **Piecewise linear** | 1363.7 | 0 |
| Linear | 1376.7 | 7.0 |
| Constant | 1581.1 | 217.4 |
| $\vert \alpha\vert $–initial velocity after stage 1 turn relationship | | |
| **Piecewise linear** | –218.1 | 0 |
| Linear | –205.1 | 13.0 |
| Constant | –171.5 | 46.6 |

$|\alpha|$ , absolute value of the turn angle; $T_1$ , time required for a displacement of 15 mm from the initial position. The best models are shown in bold.

**Table 3.** Comparison of the distribution of escape trajectories (ETs) between the model prediction (n=264 per simulation ×1000 times) and experimental data (n=264) using the two-sample Kuiper test

| Model | Median Kuiper's $V$ | Median $P$ | Rate of $P > 0.05$ |
| --- | --- | --- | --- |
| With both $D_{\text{attack}}$ and $T_1(\vert \alpha\vert )$ | 0.11 | 0.44 | 0.97 |
| With $D_{\text{attack}}$ and without $T_1(\vert \alpha\vert )$ | 0.26 | < 0.01 | 0.00 |
| Without $D_{\text{attack}}$ and with $T_1(\vert \alpha\vert )$ | 0.18 | < 0.01 | 0.12 |
| Neither $D_{\text{attack}}$ nor $T_1(\vert \alpha\vert )$ | 0.28 | < 0.01 | 0.00 |

$D_{\text{attack}}$ , distance between the prey's initial position and the endpoint of the predator attack; $T_1(|\alpha|)$ , relationship between the absolute value of the turn angle and the time required for a 15-mm displacement from the initial position (i.e., the time required for the prey to turn).

The relationship between |*α*| and the time required for a displacement of 15 mm, *T*_1_(|*α*|), is shown in Fig. 3. The time was constant up to 44° of |*α*| above which the time linearly increased in response to the increase of |*α*| (Fig. 3A). In the hierarchical Bayesian model, the lowest widely applicable or Watanabe-Akaike information criterion (WAIC) was obtained for the piecewise linear regression model (Table 2). To understand the possible mechanism of the relationship, the relationship between |*α*| and initial velocity after a stage 1 turn, calculated as the displacement per second during the 10 milliseconds (ms) after the turn, was also evaluated (Fig. 3B). The velocity increased in response to |*α*| up to 46°, beyond which it plateaus. In the hierarchical Bayesian model, the lowest WAIC was obtained for the piecewise linear regression model (Table 2). In both relationships, the regression lines by the piecewise linear model were similar to those by the GAMM, suggesting that the general trends of the relationships were clearly captured by this method. The change points of the two relationships were not significantly different [difference: 1.70±18.01° (mean±95% Bayesian credible intervals)]. These results indicate that fish with a small |*α*| (<<45°) can accomplish the stage 1 turn quickly but their velocity after the turn is lower, while fish with an intermediate |*α*| (=45°) spend a longer time on the stage 1 turn, but their velocity after the turn is higher. Fish with a large |*α*| (>> 45°) spend a still longer time on the stage 1 turn, but their velocity after the turn is similar to that with an intermediate |*α*| (Fig. 3).

We have optimized the values of *U*_pred_ and *D*_attack_ from the perspective of the prey using the experimental data (See Materials and Methods for details). Briefly, the optimal values for prey were obtained using the ranking index, where 0 means that the real fish chose the theoretically optimal ET where *T*_diff_ is the maximum, and 1 means that the real fish chose the theoretically worst ET where *T*_diff_ is the minimum (e.g., going toward the predator). The result shows that the optimal value of *D*_attack_ is 34.73 mm and the optimal value of *U*_pred_ is 1.54 m s^−1^. Using data from previously published predator-prey experiments on the same species of prey and predator [12], we show that the estimated *D*_attack_ value is at the upper limit of the empirical data and the estimated *U*_pred_ value is higher than the mean of the observed predator speed (Fig. S6). These results suggest that the values independently estimated in the present study are reasonable, and the prey may choose less risky ETs by overestimating the values of *D*_attack_ and *U*_pred_.

### Comparison of Model Predictions and Experimental Data

Figure 4A plots the relationships between the ET and the relative time difference *T*_diff_ for different initial orientations *β* estimated by the geometric model; Fig. 4B plots the relationship between the initial orientation and the theoretical ET. Forty-one percent, 76%, and 94% of observed ETs were within the top 10%, 25%, and 40% quantiles, respectively (0.1, 0.25, 0.40 ranking index) of the theoretical ETs (Figs. 4B and S7). In general, the predicted ETs are in line with the observed ones, where the model predicts a multimodal pattern of ET with a higher peak (i.e., optimal ET) at the maximum *T*_diff_ (*T*_diff,1_) and a second lower peak (i.e., suboptimal ET) at the second local maximum of *T*_diff_ (*T*_diff,2_). When the initial orientation is <20° (Figs. 4A; *β* =15°, 4B and 5B), the optimal and suboptimal ETs are around 100° (away response) and −100° (toward response), respectively, which is consistent with the bimodal distribution of our experiment (Fig. 2A; 0°≤*β*<30°). At initial orientations in the range 20‒60°, the suboptimal ET switches from around −100° to 170° (Figs. 4A; *β* =45°, 4B and 5B), although *T*_diff,2_ is extremely small compared to *T*_diff,1_ (Figs. 4A; *β* =45°, 4B and 5B). Accordingly, the second peak (i.e., at around 170°) was negligible in our experimental data (Fig. 2A; 30°≤*β* <60°), even though the fish can potentially reach such an ET (i.e., from such an initial orientation, an 170° ET is within the upper limit of |*α*|, 147°). When the initial orientation is 60‒120° (Figs. 4A; *β* =75° and *β* =105°, 4B and 5B), the optimal ET is 100‒140° (gradually shifting from 100° to 140°), and the suboptimal ET is around 170°. These two peaks and the shift of the optimal ET are consistent with the experimental results (Fig. 2A; 60°≤*β*<90° and 90°≤*β*<120°). The values of the optimal and suboptimal ETs are reversed at initial orientations >120° (Figs. 4B and 5B), as the optimal and suboptimal values become 170‒180° and around 140°, respectively (Fig. 4A). These results are again consistent with the bimodal distribution of our experiments (Fig. 2A; 120°≤*β*<150° and 150°≤*β*≤180°).

**Fig. 4.**
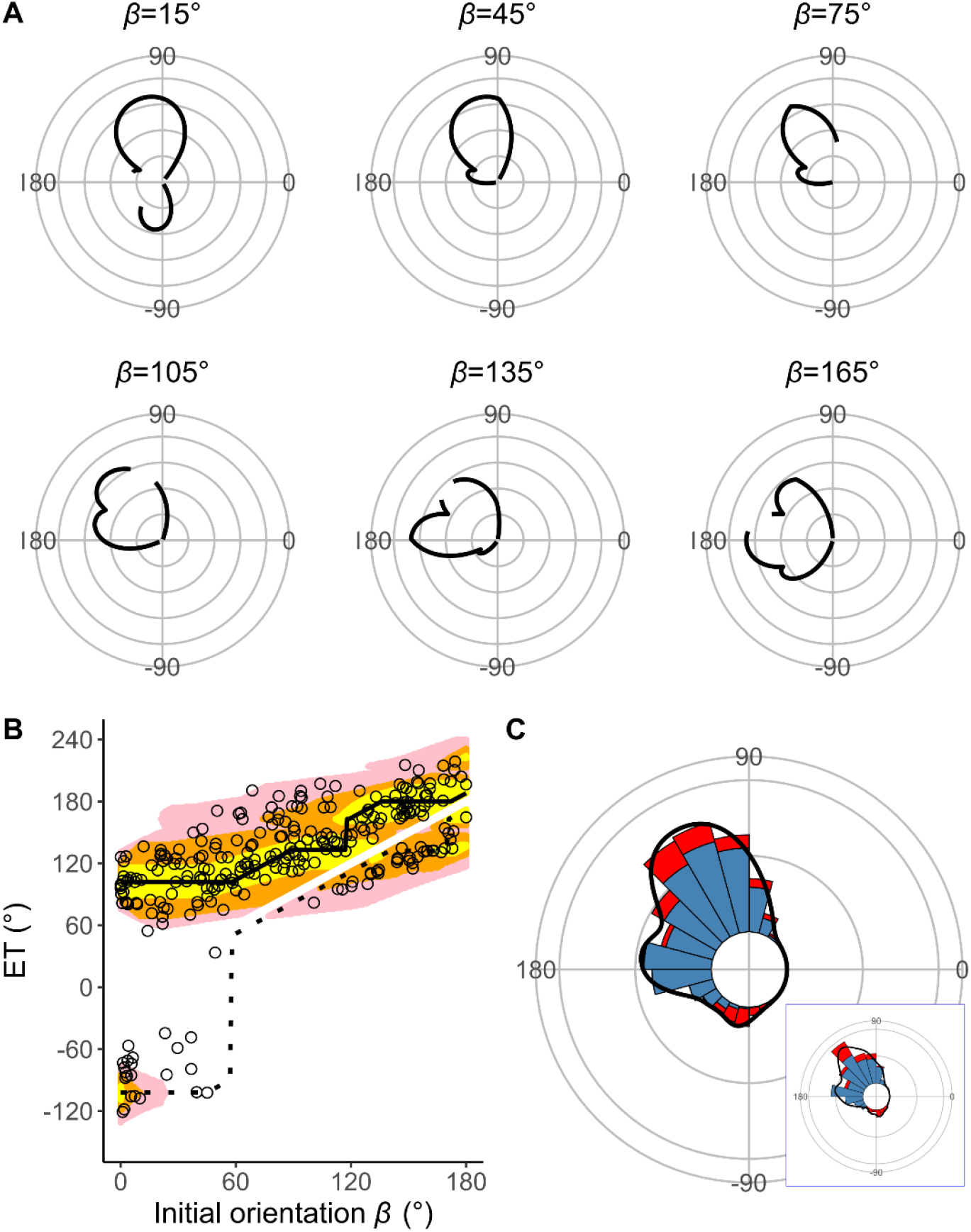
Model estimates. (A) Circular plots of the time difference between the prey and predator *T*_diff_ in different initial orientations *β* The time difference of the best escape trajectory (ET) was regarded as 10 ms, and the relative time differences between 0 and 10 ms are shown by solid lines. Areas without solid lines indicate that either the time difference is below 0 or the fish cannot reach that ET because of the constraint on the possible range of turn angles |*α*| Concentric circles represent 3 ms. (B) Relationship between the initial orientation *β* and ET. Solid and dotted lines represent the best-estimated away and toward responses, respectively. Different colors represent the top 10%, 25%, and 40% quantiles of the time difference between the prey and predator within all possible ETs. (C) Circular histogram of the theoretical ETs, estimated by a Monte Carlo simulation. The probability of selection of an ET was determined by the truncated normal distribution of the optimal ranking index (Fig. S7). This process was repeated 1000 times to estimate the frequency distribution of the theoretical ETs. Colors in the bars represent the away (blue) or toward (red) responses. Black lines represent the kernel probability density function. Concentric circles represent 10 % of the total sample sizes, the bin intervals are 15°, and the bandwidths of the kernel are 50. Circular histogram of the observed ETs (Fig. 2C) is shown in the lower right panel for comparison. The predator’s approach direction is represented by 0° (n=264 from 23 individuals for experimental data, and n=264000 for Monte Carlo simulation).

**Fig. 5.**
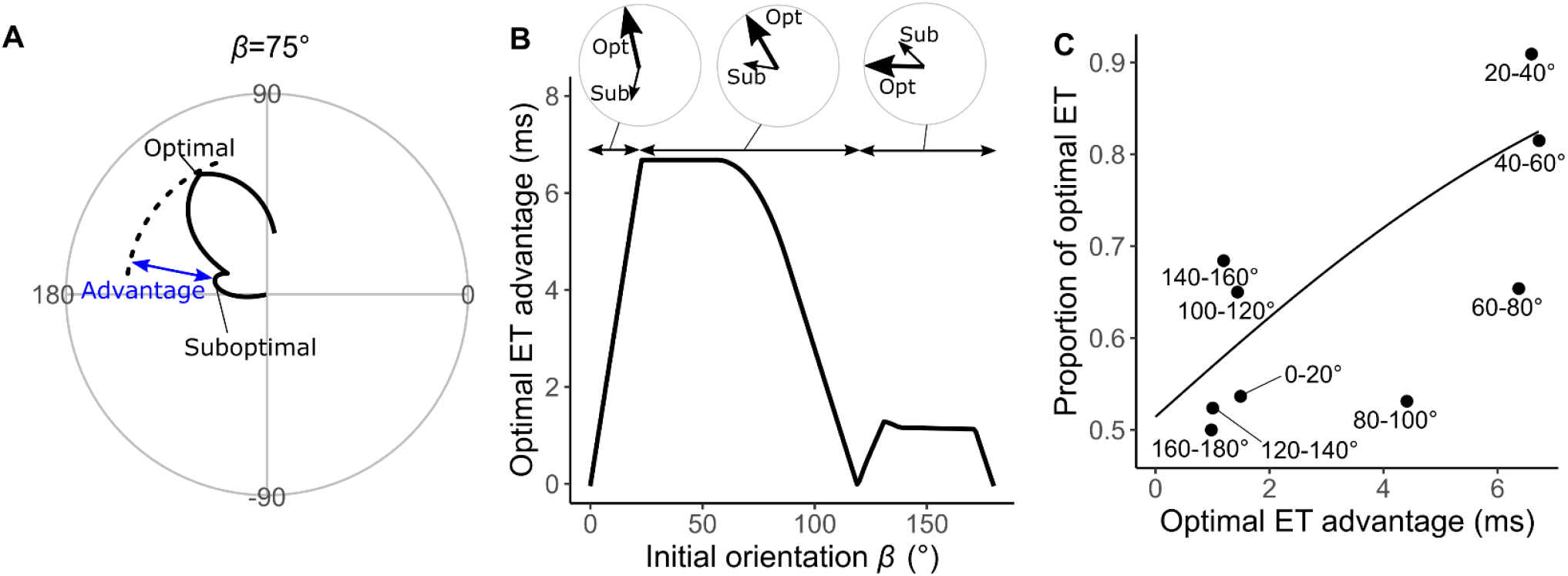
Analyses of the probability that the prey chooses the optimal vs. suboptimal ETs. (A) The time difference between the prey and predator *T*_diff_ at the initial orientation *β* of 75° is shown as an example. We defined the difference between the maximum of *T*_diff_ (at the optimal ET) and the second local maximum of *T*_diff_ (at the suboptimal ET) as the optimal ET advantage. (B) Relationship between the initial orientation *β* and the optimal ET advantage. Large and small arrows in circles represent the optimal and suboptimal ETs, respectively, for each *β* sectors. (C) Relationship between the optimal ET advantage and the proportion of the optimal ET used by the real prey in 20° initial orientation *β* bins. The line was estimated by the mixed effects logistic regression analysis (n=247 from 23 individuals).

Figure 4C shows the circular histogram of the overall theoretical ETs estimated by Monte Carlo simulation. The theoretical ETs show two large peaks at around 110–130° and 170–180°, and one small peak at around −100° (Fig. 4C). This theoretically estimated ET distribution is similar to the frequency distribution of the observed ETs (Fig. 2C); there were no significant differences in the frequency distribution between theoretical ETs (n=264 per simulation) and observed ETs (n=264) in 971 of 1000 simulations (Table 3; two-sample Kuiper test, median *V*=0.11, median *P*=0.44).

To investigate how the initial orientation of the prey modulates the proportion of using the theoretically optimal ET (i.e., where *T*_diff_ is the maximum, *T*_diff,1_) compared to using the suboptimal ET (i.e., where *T*_diff_ is the second local maximum, *T*_diff,2_), we calculated the optimal ET advantage (*T*_diff,1_−*T*_diff,2_) (Fig. 5A), which represents the difference in the buffer time available for the prey to escape from the predator, at different initial orientations. The fish chose the optimal and suboptimal ETs to a similar extent when the optimal ET advantage is negligible (Fig. 5C). For example, when looking at the optimal ET advantage <2 ms, where the initial orientation is 0‒7° and 106–180° (46% of all initial orientations), the proportion of the optimal ET used was only 55% (Fig. 5B and C). On the other hand, the proportion of the optimal ET used was 81% when the optimal ET advantage is higher than 6 ms (i.e., when the initial orientation is 21–75°) (Fig. 5B and C). There was a significant effect of optimal ET advantage on the proportion of the optimal ET used by fish tested in our experiments (Mixed-effects logistic regression analysis, *χ*^2^ =10.72, *P*<0.01, n=247).

To investigate the effects of two factors [i.e., the endpoint of the predator attack *D*_attack_ and the time required for the prey to turn *T*_1_(|*α*|)] on the predictions of ET separately, we constructed three additional geometric models (Figs. S8–S10): a model that includes only *D*_attack_, a model that includes only *T*_1_(|*α*|), and a null model that includes neither factors (Fig. 1A and [15]). In all of these models, the theoretical ET distributions estimated through Monte Carlo simulations were significantly different from the observed ET distributions (Table 3; two-sample Kuiper test, median *P*<0.01). Although the model with *D*_attack_ and the model with *T*_1_(|*α*|) show multimodal patterns of ET distribution, the simulation based on these models do not match the experimental data, likely because of differences in the values and relative heights of the peaks (Figs. S8 and S9). The null model shows a unimodal pattern of ET distribution (Fig. S10).

### Application of the model to other ET patterns

Although many fish species and animals from other taxa exhibit multiple preferred ETs similar to what we observed here, some animals show different patterns of ETs: e.g., a single preferred ET either at around 180° [42] or at around 90° [21], and multiple preferred ETs at small and large angles from the predator’s approach direction [43–45] (Fig. 6A–C). To test the hypothesis that these different ET patterns are based on the same principle of our geometric model, we changed the values of model parameters (e.g., *U*_pred_, *D*_attack_) within a realistic range, and explored whether such adjustments can produce the ET patterns observed in the original work. At small *U*_pred_, the model predicts one strong peak at around 180° (Fig. 6D), whereas at large *U*_pred_, the model predicts a strong peak at around 90° (Fig. 6E). The model where the predator can adjust the approach path and its attack lasts for a long distance (i.e., large *D*_attack_) predicts multiple preferred ETs directed at small (at around 30°) and large (at around 170°) angles from the predator’s approach direction (Fig. 6F). These results indicate that our model is flexible enough to explain various patterns of observed animal escape trajectories. See Figs. S11–S19 for details of the effect of each parameter on the ET distribution.

**Fig. 6.**
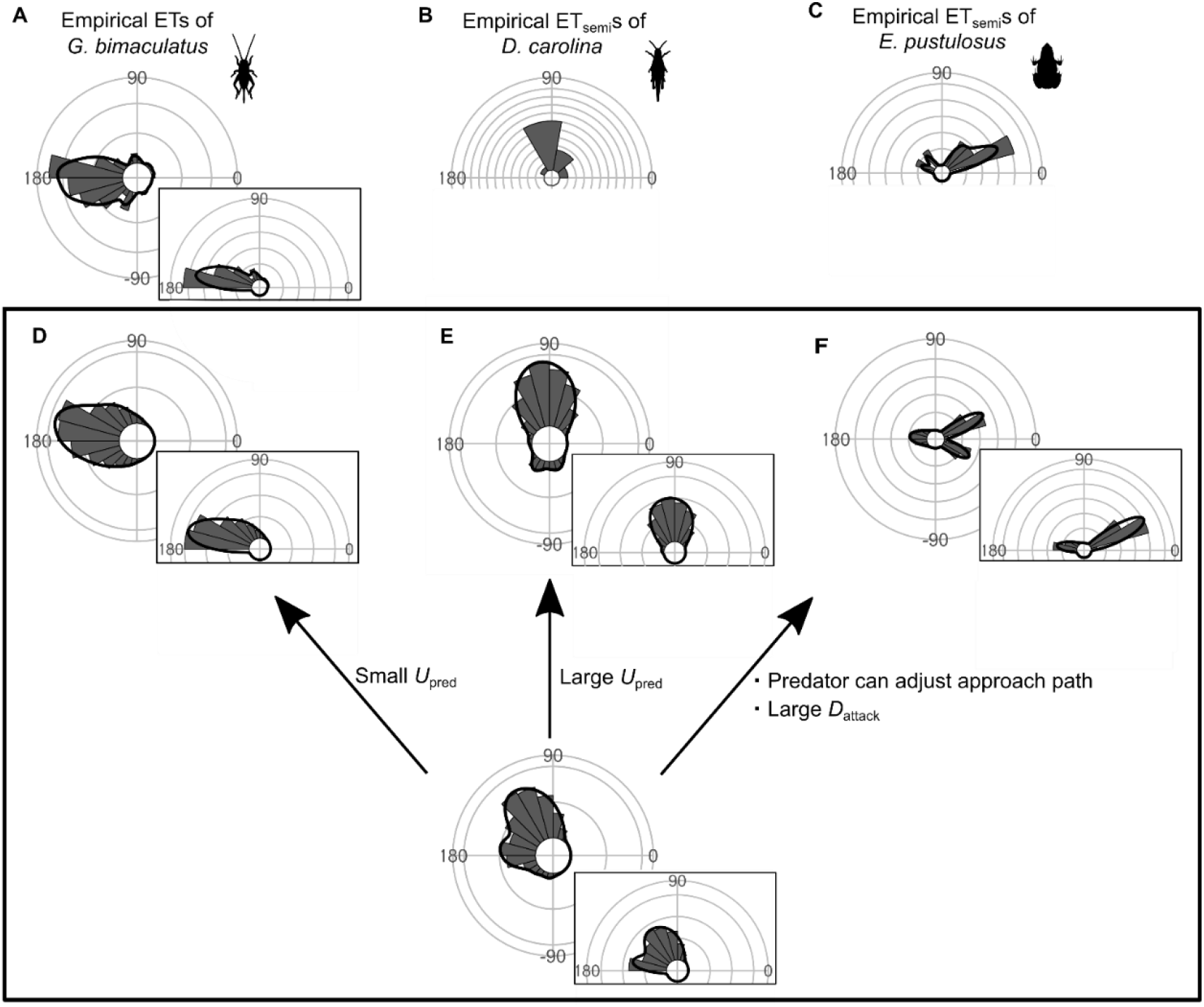
Circular histograms of other typical empirical ET distribution patterns and the potential explanations by the geometric model. Some previous studies have used the different definition for calculating the angles for escape trajectories, in which the values range from 0° (directly toward the threat) to 180° (opposite to the threat), thereby using only one semicircle regardless of their turning direction and magnitude (e.g., both 120° and 240° of ETs are regarded as 120°). This angle is denoted as ET_semi_, and is shown by a semicircular plot. (A) Unimodal ET distribution pattern at around 180° in two-spotted cricket *Gryllus bimaculatus* escaping from the air-puff stimulus. Data were obtained from Fig. 3 in Kanou et al. (1999) [42]. (B) Unimodal ET_semi_ distribution pattern at around 90° in Carolina grasshopper *Dissosteira carolina* escaping from an approaching human. Data were obtained from Fig. 2 in Cooper (2006) [21]. (C) Bimodal ET_semi_ distribution pattern directed at small and large angles from the predator’s approach direction in túngara frog *Engystomops pustulosus* escaping from an approaching dummy bat. Data were obtained from Fig. 4b in Bulbert et al. (2015) [45]. (D) Unimodal ET distribution pattern at around 180°, estimated by a Monte Carlo simulation of the geometric model. In this cases, the predator speed *U*_pred_ is very small (i.e., *K*=*U*_pred_/*U*_prey_=0.3), and the other parameter values are the same as the values used to explain the escape response of *Pagrus major* (E) Unimodal ET distribution pattern at around 90°, estimated by a Monte Carlo simulation of the model. In this case, *U*_pred_ is very large (i.e., *K*=*U*_pred_/*U*_prey_=7.5), and the other parameter values are the same as the values used to explain the escape response of *P. major* (F) Bimodal ET distribution pattern directed at small and large angles from the predator’s approach direction, estimated by a Monte Carlo simulation of the geometric model where the predator can adjust its approach path. In this case, *D*_initial_ is 130 mm, *D*_react_ is 70 mm, *R*_turn_ is 12 mm, *D*_attack_ is 400 mm, *SD*_choice_ is 0.23, and the other parameter values are the same as the values used for explaining the escape response of *P. major* Black lines represent the kernel probability density function with a bandwidth of 50, and concentric circles represent 10 % of the total sample sizes. See Table 1 and SI Appendix for details of the definitions of the variables.

## Discussion

Our geometric model, incorporating the endpoint of the predator attack, *D*_attack_, and the time required for the prey to turn, *T*_1_(|*α*|), to maximize the difference between the prey and the predator in the time of arrival at the edge of the safety zone, *T*_diff_, clearly explains the multimodal patterns of ETs in *P. major* Figure 7 shows an example of how multiple ETs result in successful escapes from predators. Specifically, according to the model, when the prey escapes at 140° or 170°, it will not be captured by the predator. On the other hand, when the prey escapes along an intermediate trajectory (157°), it will be captured because it swims toward the corner of the danger zone to exit it, and therefore it needs to travel a longer distance than when escaping at 140° or 170°. This example illustrates that the multimodal patterns of ETs are likely to be attributable to the existence of two escape routes: either moving sideways to depart from the predator’s strike path or moving opposite to the predator’s direction to outrun it. Interestingly, both components of the predator-prey interaction [i.e., *D*_attack_ and *T*_1_(|*α*|)] added to the previous model [15] are important for accurate predictions of the ET distribution because when they are considered by the model separately, the predictions do not match the experimental data (Figs. S8 and S9; Table 3).

**Fig. 7.**
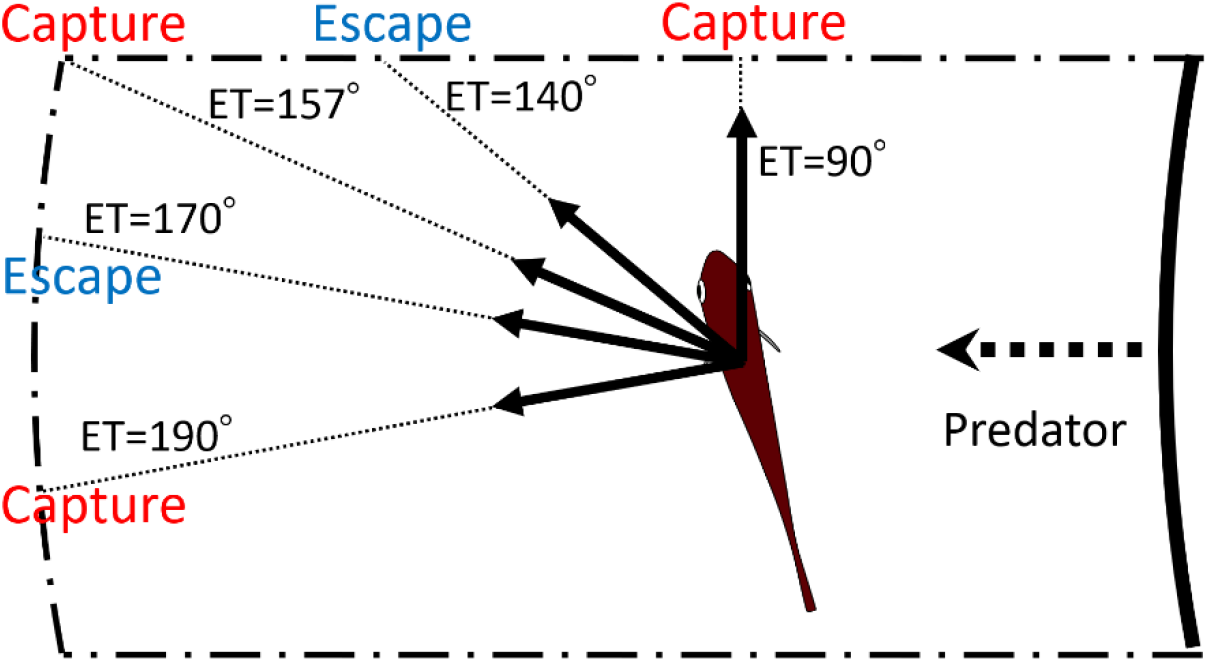
Schematic drawing showing how multiple escape trajectories (ETs) result in successful escapes from predators. The area enclosed by dash-dotted lines represents the danger zone the prey needs to exit in order to escape predation, outside of which is the safety zone. When the prey escapes toward the corner of the danger zone (ET=157°) to exit it, it needs to travel a relatively long distance and therefore the predator can catch it. On the other hand, when the prey escapes with an ET at 170° or 140°, it covers a shorter distance and can reach the safety zone before the predator’s arrival. When the prey escapes with an even smaller ET (90°), it will be captured because the shorter travel distance for the predator overrides the benefits of the smaller turn and shorter travel distance for the prey. When the prey escapes with an even larger ET (190°), it will also be captured, because the prey requires a longer time to turn than if escaping along the 170° ET, whereas the travel distance for both predator and prey is the same as that for the 170° ET. In this example, the initial orientation, flight initiation distance, and the body length posterior to the center of mass were set as 110°, 60 mm and 30 mm, respectively.

Two different escape tactics have been proposed to enhance the success of predator evasion: the optimal tactic, which maximizes *T*_diff_ (i.e., the distance between the prey and the predator) [4, 14, 15], and the protean tactic, which maximizes unpredictability to prevent predators from adjusting their strike trajectories accordingly [23–25, 46]. Our results suggest that the prey combines these two different tactics by using multiple preferred ETs. Specifically, when the optimal ET advantage is large (i.e., when the initial orientation is 20–60°), the prey mainly uses the optimal ET (Figs. 2A and 5). However, when the optimal ET advantage over the suboptimal ET is negligible (i.e., the initial orientation is close to 0° or within the range 110‒180°), the prey uses optimal and suboptimal ETs to a similar extent (Figs. 2A and 5). In such cases, the escape trajectory of the prey would be highly unpredictable for the predator. While the unpredictability at initial orientations near 0° and 180° can be easily explained by the left-right indecision at orientations nearly perpendicular to the threat [22, 40, 47], yielding ETs that are approximately symmetrical to the axis of the predator attack, the unpredictability observed at initial orientations near 110–180° is related to the similarly advantageous choice between escaping with an ET at around 140° or 180°. Interestingly, at initial orientations >120°, our results show that these two ETs are reached by using toward and away responses, respectively. The overlap between the ETs of toward and away responses in the overall dataset (Fig. 2) suggests that toward responses are not “tactical mistakes” of the prey that turns toward a threat, but are simply related to reaching an optimal or suboptimal ET. These results suggest that the prey strategically adjusts the use of optimal and protean tactics based on their initial orientation. This allows the prey to have unpredictable ETs, thereby preventing predators from anticipating their escape behavior, while keeping *T*_diff_ large enough to enter the safety zone before the predator reaches it.

A relevant question from a perspective of neurosensory physiology is how the animals are able to determine their ETs within milliseconds of response time. The initial orientation of the prey has been incorporated into various neural circuit models [48–51], but these models assume that prey animals always escape in a 180° direction (i.e., opposite to the stimulus source), irrespective of the initial orientation. However, the present study shows that animals use suboptimal ETs as well as optimal ETs, and that these ETs may change in a nonlinear fashion, depending on the initial orientation. Thus, we require new neurophysiological models of ETs to understand how neural circuits process the sensory cues of a threatening stimulus, resulting in muscle actions that generate multiple preferred ETs.

Our geometric model assumes that the prey determines the ETs based on a fixed predator speed. This assumption is supported by the results of our experiments, where the effects of predator speed on the mean and variability of ETs are not significant. Although we did not find any effect of predator speed, it is possible that a speed outside the range we used may affect ETs. Recent studies show that larval zebrafish *Danio rerio* exhibit less variable ETs under faster threats than they do under slower threats [52, 53], and the difference in ET variability between fast and slow threats is dependent on whether the Mauthner cell is active or not [53]. Therefore, any differences in the ET variability of the present study compared to previous studies could be related to the different involvement of the Mauthner-cells. Using the conventional geometric model, Soto *et al.* [14] showed that the choice of ET only matters to a prey when the predator speed is intermediate, because a prey that is much faster than its predator can escape by a broad range of ETs, whereas a prey that is much slower than its predator cannot escape by any ETs [16]. The predator speed used in this study is in the range of the real predator speed in the previous study using the same species of both predator and prey [12]. Thus, our results are ecologically relevant, and the prey is likely to have optimized their ETs based on a fixed predator speed, where the choice of ET strongly affects their survival.

The relationship between |*α*| and the time required for a 15-mm displacement, *T*_1_(|*α*|), (Fig. 3A) indicates that the time required for a 15-mm displacement is relatively constant up to an |*α*| of about 45°, while a further change in |*α*| requires additional time. This relationship is likely to be attributable to the kinematics and hydrodynamics of the C-start escape response, because the initial velocity after the stage 1 turn increases linearly up to about 45°, beyond which it plateaus (Fig. 3B). Interestingly, a recent study on swimming efficiency during acceleration found that efficiency increases linearly with yaw amplitudes up to a certain value, beyond which efficiency plateaus [54].

We show that our model can explain other empirically observed ET patterns (Figs. 6, S11–S19). Based on the model assuming that the predator makes an in-line attack toward the prey, which is realistic for ambush and stalk-and-attack predators [55] [e.g., frogs [11], spiders [13], and fish [12, 28, 29, 56]], either single or multiple ETs at around 90–150° and around 180° are predicted, as have been observed in many empirical studies of animals escaping from ambush predators and artificial stimuli [20]. Based on the model assuming that the predator can adjust its approach path, which is realistic for pursuit predators, multiple ETs directed at small and large angles from the predator’s approach direction can be predicted, as observed in the empirical studies of prey escaping from pursuit predators [18, 45]. These results suggest that our model is flexible enough to be applied to various predator-prey systems.

Our work represents a major advancement in understanding the basis of the variability in ETs observed in previous works (reviewed in [20]). Our results suggest that prey use multiple preferred ETs to maximize the time difference between themselves and the attacking predator, while keeping a high level of unpredictability. The results also suggest that prey strategically adjust the use of protean and optimal tactics with respect to the advantage of the optimal ET over the suboptimal ET. Because multimodal ETs similar to what we observed here have been found in many fish species and other animal taxa [20], this behavioral phenotype may result from convergent evolution in phylogenetically distant animals. From a neurosensory perspective, our findings open new avenues to investigate how the animals determine their ETs from multiple options with specific probabilities, which are modulated by the initial orientation with respect to the threat.

## Materials and Methods

### Definition of the Angles

The C-start escape response consists of an initial bend (stage 1), followed by a return tail flip (stage 2), and continuous swimming or coasting (stage 3) [37, 38]. In line with previous studies [20, 40, 57], we defined directionality (away or toward responses), initial orientation *β* turn angle *α* and ET *α+β* as follows (Fig. S1). *Directionality* the away and toward responses were defined by the first detectable movement of the fish in a direction either away from or toward the predator, respectively [20]. *Initial orientation* (*β*): the angle between the line passing through the prey’s center of mass [CoM; located at 34% of the total length from the tip of the snout [12]] and the tip of the snout at the onset of stage 1, and the midline of the predator model attacking in a straight line. Initial orientation ranges from 0° (i.e., when the prey is attacked from front) to 180° (i.e., when the prey is attacked from behind). *Turn angle* (*α*): the angle between the line passing through the CoM and the tip of the snout at the onset of stage 1, and the line passing through the CoM at the onset of stage 1 and the CoM at the end of stage 2. The angles of the away and toward responses are assigned positive and negative values, respectively. *ET* (*α+β*): the sum of the initial orientation (*β*) and the turn angle (*α*). ET is a circular variable since it can span 360°. Because the experimental data exhibited no asymmetry in directionality (Fisher’s exact test, *P*=1.00, n=264) and ET distribution (two-sample Kuiper test, *V*=0.14, *P*=0.61, n=264), we pooled the ETs from the left and right sides, treating all fish as though they were attacked from the right side [20].

### Experiment

We have elicited the escape response of *P. major* [45.33±3.48 mm (mean±s.d.) total length, n=23] using a dummy predator. The experiment was conducted in a plastic tank (540×890×200 mm) filled with seawater to a depth of 80 mm. The water temperature was maintained at 23.8 to 24.7℃. An individual *P. major* was introduced into a PVC pipe (60 mm diameter) set in the center of the tank and acclimated for 15 min. After the acclimation period, the PVC pipe was slowly removed, and the dummy predator, a cast of *Sebastiscus marmoratus* (164 mm in total length and 36 mm in mouth width), was moved toward the *P. major* for a distance of 200 mm by using a plastic rubber band (Fig. S20). Because the previous work shows that *S. marmoratus* attacks *P. major* using a variable speed [1.10±0.65 (0.09–2.31) m s^−1^, mean±s.d. (range)] [12], we used various strengths of plastic rubber bands to investigate the effect of predator speed on ET. The fish movements were recorded from above, using a high-speed video camera (HAS-L1; Ditect Co., Tokyo, Japan) at 500 frames s^−1^. Each individual *P. major* was recorded from 5 to 20 times, and, in total, we obtained 264 escape response data. The recorded videos were analyzed frame by frame using Dipp-Motion Pro 2D (Ditect Co.). The CoM and the tip of the mouth of *P. major* and the tip of the predator’s mouth were digitized in each frame to calculate all the kinematic variables. The animal care and experimental procedures were approved by the Animal Care and Use Committee of the Faculty of Fisheries (Permit No. NF-0002), Nagasaki University in accordance with the Guidelines for Animal Experimentation of the Faculty of Fisheries and the Regulations of the Animal Care and Use Committee, Nagasaki University.

Because our geometric model predicts that the initial orientation *β* and the predator speed *U*_pred_ affect the ET and turn angle *α* we examined these effects by the experimental data using a GAMM with a normal distribution and identity link function [58]. ET and *α* were regarded as objective variables, while predator speed and initial orientation were regarded as explanatory variables and were modeled with a B-spline smoother. Fish ID was regarded as a random factor. Smoothed terms were fitted using penalized regression splines, and the amount of smoothing was determined using the restricted maximum likelihood (REML) method. As was done in previous studies [20, 39, 40], the away and toward responses were analyzed separately. The significance of the initial orientation and predator speed was assessed by the *F*-test. The analysis was conducted using R 3.5.3 (R Foundation for Statistical Computing) with the R package *gamm4*.

### Determination of Parameter Values

#### Determination of the Prey’s Kinematic Parameters

The relationship between |*α*| and the time required for a displacement of 15 mm, *T*_1_(|*α*|), was estimated by piecewise linear regression [59]. We used piecewise linear regression rather than a commonly used smoothing method such as GAMM, because the smoothing method does not output the timing of the regression change and thus the biological interpretation of the regression curve is problematic [59]. The time required for a displacement of 15 mm was regarded as an objective variable, whereas |*α*| was regarded as an explanatory variable. Fish ID was included as a covariate in order to take into account potential individual differences in the relationship, *T*_1_(|*α*|). To detect the possible kinematic mechanism of the relationship *T*_1_(|*α*|), we also examined the relationship between |*α*| and initial velocity after the stage 1 turn, using piecewise linear regression. Initial velocity after the stage 1 turn was regarded as an objective variable, |*α*| was regarded as an explanatory variable, and fish ID was included as a covariate. A hierarchical Bayesian model with a Markov chain Monte Carlo (MCMC) method was used to estimate these relationships [59, 60]. The number of draws per chain, thinning rate, burn-in length, and number of chains were set as 200000, 1, 100000, and 5, respectively. To test the overall fit of the model, the WAIC of the model was compared with those of the null model (constant) and a simple linear regression model. MCMC was conducted using RStan 2.18.2 (Stan Development Team 2019).

#### Determination of Predator speed and Endpoint of the Predator Attack

Because we had no previous knowledge about the values of *U*_pred_ and *D*_attack_ that the prey regards as dangerous (i.e., the values of *U*_pred_ and *D*_attack_ that trigger a response in the prey), we optimized the values using the experimental data in this study. We have input the obtained values of *D*_width_, *R*_device_, *D*_1_, *U*_prey_, and *T*_1_(|*α*|) into the theoretical model. The optimal values of *U*_pred_ and *D*_attack_ were obtained using the ranking index. The ranks of the observed ETs among the theoretical ET choices of 1° increment were standardized as the ranking index, where 0 means that the real fish chose the theoretically optimal ET where *T*_diff_ is the maximum, and 1 means that the real fish chose the theoretically worst ET where *T*_diff_ is the minimum. The optimal set of *D*_attack_ and *U*_pred_ values was determined by minimizing the mean ranking index of the observed ETs. The distribution of the optimal ranking index was then fitted to the truncated normal distribution and was used to predict how the fish chose the ETs from the continuum of the theoretically optimal and worst ETs.

### Model Predictions

We input the above parameters [*D*_width_, *R*_device_, *D*_1_, *U*_prey_, *T*_1_(|*α*|), *D*_attack_, and *U*_pred_] into the model and calculated how the choice of different ETs affects *T*_diff_ for each initial orientation *β* Because there was a constraint on the possible range of |*α*| (i.e., fish escaping by C-start have a minimum and maximum |*α*| [33]), the range of |*α*| was determined based on its minimum and maximum values observed in our experiment, which were 9~147°.

To estimate the overall frequency distribution of ETs that include the data on observed initial orientations, we conducted Monte Carlo simulations. In each observed initial orientation, the ET was chosen from the continuum of the theoretically optimal and worst ETs. The probability of the ET selection was determined by the truncated normal distribution of the optimal ranking index (e.g., the fish could choose theoretically good ETs with higher probability than theoretically bad ETs, but the choice is a continuum based on the truncated normal distribution). This process was repeated 1000 times to robustly estimate the frequency distribution of the theoretical ETs. In each simulation run, the frequency distribution of the theoretical ETs was compared with that of the observed ETs using the two-sample Kuiper test [61].

To investigate how the real prey changes the probability that it uses the theoretically optimal ET or suboptimal ET, we regarded the difference between the maximum of *T*_diff_ (at the optimal ET) and the second local maximum of *T*_diff_ (at the suboptimal ET) as the optimal ET advantage, and theoretically estimated the values for all initial orientations. We then examined the relationship between the optimal ET advantage and the proportion of the optimal ET the prey actually chose using a mixed-effects logistic regression analysis [58]. Each observed ET was designated as the optimal (1) or the suboptimal (0) based on whether the observed ET was closer to the optimal ET or suboptimal ET. When the prey chose the ET that was more than 35° different from both the optimal and suboptimal ETs, the ET data point was removed from the analysis (these cases were rare: 7%). The choice of ET [optimal (1) or suboptimal (0)] was regarded as an objective variable, while the optimal ET advantage was regarded as an explanatory variable. Fish ID was regarded as a random factor. The significance of the optimal ET advantage was assessed by the likelihood ratio test with *χ*^2^ distribution. The analysis was conducted using R 3.5.3 with the R package *lme4*.

To investigate the effects of two factors [i.e., the endpoint of the predator attack *D*_attack_ and the time required for the prey to turn *T*_1_(|*α*|)] on predictions of ET separately, we compared four geometric models: the model that includes both *D*_attack_ and *T*_1_(|*α*|), the model that includes only *D*_attack_, the model that includes only *T*_1_(|*α*|), and the null model. Note that the null model is equivalent to the previous Domenici’s model [15]. In all models, the values of *U*_pred_ and *D*_attack_ were optimized using the ranking index. The overall frequency distributions of ETs were estimated through Monte Carlo simulations, and in each simulation run, the theoretical ET distribution was compared with the observed ET distribution using the two-sample Kuiper test.

To investigate whether our model can explain other empirical ET patterns, we changed the values of model parameters (e.g., *U*_pred_, *D*_attack_) within a realistic range, and conducted Monte Carlo simulations to estimate the frequency distribution of the theoretical ETs. For each initial orientation, ranging from 0° to 180° with an increment of 1°, the ET was chosen based on the probability of the truncated normal distribution (i.e., the continuum of the theoretically optimal and worst ETs), and this process was repeated 100 times. In the model where the predator cannot adjust the strike path (Fig. 1C), we fixed three parameters and varied the fourth parameter (*U*_pred_, *D*_attack_, *R*_device_, and s.d. of the truncated normal distribution for ET choice, *SD*_choice_) from the model produced for the escape response of *P. major* (i.e., *D*_attack_=34.73 mm, *U*_pred_=1.54 m s^−1^, *R*_device_=199 mm, *SD*_choice_=0.33). Using the model where the predator can adjust the strike path (Figs. 1D and S3), we simulate the situation in which the safety zone shape inside the predator’s turning radius is similar to the Corcoran’s model (Fig. 1B) but we included a safety zone opposite to the incoming direction of the predator. We considered *D*_attack_ as 400 mm, *D*_initial_ as 130 mm, the minimum turning radius of the predator *R*_turn_ as 12 mm, and the reaction distance of the predator *D*_react_ as 70 mm. We used the same values of the *P. major* model for *R*_device_ and the other parameters. We then fixed four parameters and varied the fifth parameter (*U*_pred_, *D*_attack_, *D*_initial_, *R*_turn,_ *SD*_choice_) to examine the effect of each parameter on the ET distribution.

## Author contributions

Y.K. conceived the study; Y.K. and P.D. constructed the theoretical model; Y.K. and H.A. designed the experiment, Y.K., H.A., H.K., and N.N. conducted the experiment; H.A. and H.K. digitized the videos; Y.K., K.S., G.N.N., and P.D. conducted statistical analyses; Y.K. wrote the manuscript with input from K.S., G.N.N., N.N., and P.D.

## Acknowledgements

We sincerely thank Y. Y. Watanabe for his constructive comments on an early version of this paper. We also thank H. Kamihata for providing equipment for rearing *P. major* This study was funded by Grants-in-Aid for Scientific Research, Japan Society for the Promotion of Science, to Y.K. (17K17949 and 19H04936), Sumitomo Foundation to Y.K. (153128), and the ISM Cooperative Research Program to Y.K. and K.S (2014-ISM.CRP-2006).

## References

1. Bullock TH. Comparative neuroethology of startle, rapid escape, and giant fiber-mediated responses. In: Eaton RC, editor. Neural mechanisms of startle behavior. Boston, MA: Springer; 1984. p. 1–13.

2. Cooper WE, Blumstein DT. Escaping from predators. Cambridge, UK: Cambridge University Press; 2015. 442 p.

3. Meager JJ, Domenici P, Shingles A, Utne-Palm AC. Escape responses in juvenile Atlantic cod *Gadus morhua* L.: the effects of turbidity and predator speed. J Exp Biol. 2006;209(20):4174–84.

4. Arnott SA, Neil DM, Ansell AD. Escape trajectories of the brown shrimp *Crangon crangon* and a theoretical consideration of initial escape angles from predators. J Exp Biol. 1999;202(2):193–209.

5. Bateman PW, Fleming PA. Switching to Plan B: Changes in the escape tactics of two grasshopper species (Acrididae: Orthoptera) in response to repeated predatory approaches. Behav Ecol Sociobiol. 2014;68(3):457–65. doi: 10.1007/s00265-013-1660-0.

6. Hein AM, Gil MA, Twomey CR, Couzin ID, Levin SA. Conserved behavioral circuits govern high-speed decision-making in wild fish shoals. Proc Natl Acad Sci USA. 2018;115(48):12224–8. doi: 10.1073/pnas.1809140115.

7. Broom M, Ruxton GD. You can run or you can hide: Optimal strategies for cryptic prey against pursuit predators. Behav Ecol. 2005;16(3):534–40. doi: 10.1093/beheco/ari024.

8. Cooper WE, Pérez-Mellado V, Baird T, Baird TA, Caldwell JP, Vitt LJ. Effects of risk, cost, and their interaction on optimal escape by nonrefuging Bonaire whiptail lizards, *Cnemidophorus murinus*. Behav Ecol. 2003;14(2):288–93. doi: 10.1093/beheco/14.2.288.

9. Walker JA, Ghalambor CK, Griset OL, McKenney D, Reznick DN. Do faster starts increase the probability of evading predators? Funct Ecol. 2005;19(5):808–15.

10. Shifferman E, Eilam D. Movement and direction of movement of a simulated prey affect the success rate in barn owl *Tyto alba* attack. J Avian Biol. 2004;35(2):111–6. doi: 10.1111/j.0908-8857.2004.03257.x.

11. Camhi JM, Tom W, Volman S. The escape behavior of the cockroach *Periplaneta americana* II. Detection of natural predators by air displacement. J Comp Physiol A Sens Neural Behav Physiol. 1978;128(3):203–12. doi: 10.1007/bf00656853.

12. Kimura H, Kawabata Y. Effect of initial body orientation on escape probability of prey fish escaping from predators. Biol Open. 2018;7(7):bio023812. doi: 10.1242/bio.023812.

13. Dangles O, Ory N, Steinmann T, Christides JP, Casas J. Spider’s attack versus cricket’s escape: velocity modes determine success. Anim Behav. 2006;72(3):603–10. doi: 10.1016/j.anbehav.2005.11.018.

14. Weihs D, Webb PW. Optimal avoidance and evasion tactics in predator-prey interactions. J Theor Biol. 1984;106(2):189–206.

15. Domenici P. The visually mediated escape response in fish: predicting prey responsiveness and the locomotor behaviour of predators and prey. Mar Freshwat Behav Physiol. 2002;35(1-2):87–110. doi: 10.1080/10236240290025635.

16. Soto A, Stewart WJ, McHenry MJ. When optimal strategy matters to prey fish. Integr Comp Biol. 2015;55(1):110–20. doi: 10.1093/icb/icv027.

17. Howland HC. Optimal strategies for predator avoidance: The relative importance of speed and manoeuvrability. J Theor Biol. 1974;47(2):333–50. doi: 10.1016/0022-5193(74)90202-1.

18. Corcoran AJ, Conner WE. How moths escape bats: predicting outcomes of predator-prey interactions. J Exp Biol. 2016;219:2704–15. doi: 10.1242/jeb.137638.

19. Martin BT, Gil MA, Fahimipour AK, Hein AM. Informational constraints on predator–prey interactions. Oikos. 2021. doi: 10.1111/oik.08143.

20. Domenici P, Blagburn JM, Bacon JP. Animal escapology II: Escape trajectory case studies. J Exp Biol. 2011;214(15):2474–94.

21. Cooper WE. Risk factors and escape strategy in the grasshopper *Dissosteira carolina*. Behaviour. 2006;143(10):1201–18. doi: 10.1163/156853906778691595.

22. Domenici P, Blake RW. Escape trajectories in angelfish (*Pterophyllum eimekei*). J Exp Biol. 1993;177:253–72.

23. Domenici P, Blagburn JM, Bacon JP. Animal escapology I: Theoretical issues and emerging trends in escape trajectories. J Exp Biol. 2011;214(15):2463–73.

24. Humphries DA, Driver PM. Protean defence by prey animals. Oecologia. 1970;5(4):285–302. doi: 10.1007/bf00815496.

25. Jones KA, Jackson AL, Ruxton GD. Prey jitters; protean behaviour in grouped prey. Behav Ecol. 2011;22(4):831–6. doi: 10.1093/beheco/arr062.

26. Richardson G, Dickinson P, Burman OHP, Pike TW. Unpredictable movement as an anti-predator strategy. Proc R Soc Lond B. 2018;285(1885). doi: 10.1098/rspb.2018.1112.

27. Moore TY, Cooper KL, Biewener AA, Vasudevan R. Unpredictability of escape trajectory explains predator evasion ability and microhabitat preference of desert rodents. Nature Communications. 2017;8(1). doi: 10.1038/s41467-017-00373-2.

28. Webb PW, Skadsen JM. Strike tactics of *Esox*. Can J Zool. 1980;58(8):1462–9.

29. Fouts WR, Nelson DR. Prey capture by the pacific angel shark, *Squatina californica* visually mediated strikes and ambush-site characteristics. Copeia. 1999;1999(2):304–12. doi: 10.2307/1447476.

30. Anderson CW. The modulation of feeding behavior in response to prey type in the frog *Rana pipiens*. J Exp Biol. 1993;179(1):1–12.

31. Paglianti A, Domenici P. The effect of size on the timing of visually mediated escape behaviour in staghorn sculpin *Leptocottus armatus*. J Fish Biol. 2006;68:1177–91. doi: 10.1111/j.1095-8649.2006.00991.x.

32. Fabian ST, Sumner ME, Wardill TJ, Rossoni S, Gonzalez-Bellido PT. Interception by two predatory fly species is explained by a proportional navigation feedback controller. J R Soc Lond Interface. 2018;15(147). doi: 10.1098/rsif.2018.0466.

33. Domenici P, Blake RW. The kinematics and performance of the escape response in the angelfish (*Pterophyllum eimekei*). J Exp Biol. 1991;156:187–205.

34. Danos N, Lauder GV. Challenging zebrafish escape responses by increasing water viscosity. J Exp Biol. 2012;215(Pt 11):1854–62. doi: 10.1242/jeb.068957.

35. Fleuren M, van Leeuwen JL, Quicazan-Rubio EM, Pieters RPM, Pollux BJA, Voesenek CJ. Three-dimensional analysis of the fast-start escape response of the least killifish, *Heterandria formosa*. J Exp Biol. 2018;221(Pt 7). doi: 10.1242/jeb.168609.

36. Ellerby DJ, Altringham JD. Spatial variation in fast muscle function of the rainbow trout *Oncorhynchus mykiss* during fast-starts and sprinting. J Exp Biol. 2001;204(13):2239–50.

37. Domenici P, Blake RW. The kinematics and performance of fish fast-start swimming. J Exp Biol. 1997;200(8):1165–78.

38. Weihs D. The mechanism of rapid starting of slender fish. Biorheology. 1973;10(3):343–50.

39. Domenici P, Booth D, Blagburn JM, Bacon JP. Escaping away from and towards a threat: The cockroach’s strategy for staying alive. Communicative and Integrative Biology. 2009;2(6):497–500.

40. Nair A, Changsing K, Stewart WJ, McHenry MJ. Fish prey change strategy with the direction of a threat. Proc R Soc Lond B. 2017;284(1857). doi: 10.1098/rspb.2017.0393.

41. Domenici P, Booth D, Blagburn JM, Bacon JP. Cockroaches keep predators guessing by using preferred escape trajectories. Curr Biol. 2008;18(22):1792–6. doi: 10.1016/j.cub.2008.09.062.

42. Kanou M, Ohshima M, Inoue J. The air-puff evoked escape behavior of the cricket *Gryllus bimaculatus* and its compensational recovery after cercal ablations. Zool Sci. 1999;16(1):71–9. doi: 10.2108/zsj.16.71.

43. Fuiman LA. Development of predator evasion in Atlantic herring, *Clupea harengus* L. Anim Behav. 1993;45(6):1101–16. doi: 10.1006/anbe.1993.1135.

44. Martín J, López P. The escape response of juvenile *Psammodromus algirus* lizards. J Comp Psychol. 1996;110(2):187–92.

45. Bulbert MW, Page RA, Bernal XE. Danger comes from all fronts: Predator-dependent escape tactics of túngara frogs. PLoS ONE. 2015;10(4):e0120546. doi: 10.1371/journal.pone.0120546.

46. Kurvers RHJM, Krause S, Viblanc PE, Herbert-Read JE, Zaslansky P, Domenici P, et al. The evolution of lateralization in group hunting sailfish. Curr Biol. 2017;27(4):521–6. doi: 10.1016/j.cub.2016.12.044.

47. Domenici P, Batty RS. Escape behaviour of solitary herring (*Clupea harengus*) and comparisons with schooling individuals. Mar Biol. 1997;128(1):29–38.

48. Eaton RC, Lee RKK, Foreman MB. The Mauthner cell and other identified neurons of the brainstem escape network of fish. Prog Neurobiol. 2001;63(4):467–85.

49. Yono O, Shimozawa T. Synchronous firing by specific pairs of cercal giant interneurons in crickets encodes wind direction. Biosystems. 2008;93(3):218–25. doi: 10.1016/j.biosystems.2008.04.014.

50. Card GM. Escape behaviors in insects. Curr Opin Neurobiol. 2012;22(2):180–6. doi: 10.1016/j.conb.2011.12.009.

51. Levi R, Camhi JM. Population vector coding by the giant interneurons of the cockroach. J Neurosci. 2000;20(10):3822–9.

52. Stewart WJ, Nair A, Jiang H, McHenry MJ. Prey fish escape by sensing the bow wave of a predator. J Exp Biol. 2014;217(24):4328–36. doi: 10.1242/jeb.111773.

53. Bhattacharyya K, McLean DL, MacIver MA. Visual threat assessment and reticulospinal encoding of calibrated responses in larval zebrafish. Curr Biol. 2017;27(18):2751–62 e6. doi: 10.1016/j.cub.2017.08.012.

54. Akanyeti O, Putney J, Yanagitsuru YR, Lauder GV, Stewart WJ, Liao JC. Accelerating fishes increase propulsive efficiency by modulating vortex ring geometry. Proc Natl Acad Sci USA. 2017;114(52):13828–33. doi: 10.1073/pnas.1705968115.

55. Moore TY, Biewener AA. Outrun or outmaneuver: predator–prey interactions as a model system for integrating biomechanical studies in a broader ecological and evolutionary context. Integr Comp Biol. 2015;55(6):1188–97. doi: 10.1093/icb/icv074.

56. Rand DM, Lauder GV. Prey capture in the chain pickerel, *Esox niger* correlations between feeding and locomotor behavior. Can J Zool. 1981;59(6):1072–8.

57. Stewart WJ, Cardenas GS, McHenry MJ. Zebrafish larvae evade predators by sensing water flow. J Exp Biol. 2013;216(3):388–98.

58. Zuur A, Ieno EN, Walker N, Saveliev AA, Smith GM. Mixed effects models and extensions in ecology with R. New York: Springer; 2009. 574 p.

59. Brilleman SL, Howe LD, Wolfe R, Tilling K. Bayesian piecewise linear mixed models with a random change point: an application to BMI rebound in childhood. Epidemiology. 2017;28(6):827–33. doi: 10.1097/EDE.0000000000000723.

60. Kéry M, Schaub M. Bayesian population analysis using winbugs – a hierarchical perspective. Burlington: Academic Press; 2011. 535 p.

61. Zar JH. Biostatistical analysis: fifth edition. New Jersey: Pearson Education; 2010. 944 p.

