## Supplementary Information for "Geometric model incorporating prey’s turn and predator attack endpoint explains multiple preferred escape trajectories"

**This PDF file includes:**

Supplementary Information text
Figures S1 to S20
Tables S1 to S3
SI References

**Supplementary Information Text**

**Mathematical formula for the geometric model modified from Corcoran & Conner (2016) [1].**

When the prey's center of mass (CoM) at the onset of its escape is located at point (0, 0), the trajectory of the CoM ($X_{\mathrm{prey}}$, $Y_{\mathrm{prey}}$) is given by:

$$Y_{\mathrm{prey}} = X_{\mathrm{prey}} \tan(\alpha + \beta) \qquad [1]$$

The edge of the safety zone is determined by the half-width of the predator capture device $D_{\mathrm{width}}$, the distance between the prey and the predator at the onset of the prey's escape response $D_{\mathrm{initial}}$, the distance required for the predator to react the prey's escape response to initiate its turn $D_{\mathrm{react}}$, the distance between the prey's initial position and the tip of the predator capture device at the end of the predator attack $D_{\mathrm{attack}}$, the minimum turning radius of the predator $R_{\mathrm{turn}}$, and the shape of the predator's capture device at the moment of attack, which is approximated as an arc with a certain radius $R_{\mathrm{device}}$. The edge of the safety zone can be divided into four parts: 1) the straight line before the onset of the predator's turn, 2) the arc of the minimum inner turning radius, 3) the capture device shape at the end of the predator attack when it attacks with the minimum turning radius, 4) the involute curve where the tip of the predator capture device traverses a specific distance from the initial position, which can be described by the trace of unwrapping a taut string from the minimum turning circle. Note that the model may lack some parts, depending on the values of parameters. The projection of the predator's capture device edge along the $X$-axis $D_2$ can be expressed as:

$$D_2 = R_{\mathrm{device}}\{1 - \cos(\sin^{-1}\frac{D_{\mathrm{width}}}{R_{\mathrm{device}}})\} \qquad [2]$$

The angle for the predator to traverse with the minimum turning radius $\gamma$ can be expresses as:

$$\gamma = \frac{D_{\mathrm{initial}} - D_{\mathrm{react}} + D_{\mathrm{attack}}}{R_{\mathrm{turn}}} \qquad [3]$$

The x and y coordinates of the change point between the 1st and 2nd parts of the safety zone edge ($X_{\mathrm{cp1}}$, $Y_{\mathrm{cp1}}$) can be expressed as:

$$\begin{cases} X_{\mathrm{cp1}} = D_{\mathrm{initial}} - D_{\mathrm{react}} + D_2 \\ Y_{\mathrm{cp1}} = D_{\mathrm{width}} \end{cases} \qquad [4]$$

The x and y coordinates of the change point between the 2nd and 3rd parts of the safety zone edge ($X_{\mathrm{cp2}}$,

$Y_{\mathrm{cp2}}$) can be expressed as:

$$\begin{cases} X_{\mathrm{cp2}} = D_2 \cos\gamma + (D_{\mathrm{width}} - R_{\mathrm{turn}}) \sin\gamma + D_{\mathrm{initial}} - D_{\mathrm{react}} \\ Y_{\mathrm{cp2}} = -D_2 \sin\gamma + (D_{\mathrm{width}} - R_{\mathrm{turn}}) \cos\gamma + R_{\mathrm{turn}} \end{cases} \quad [5]$$

The x and y coordinates of the change point between the 3rd and 4th parts of the safety zone edge ($X_{\mathrm{cp3}}$, $Y_{\mathrm{cp3}}$) can be expressed as:

$$\begin{cases} X_{\mathrm{cp3}} = -R_{\mathrm{turn}} \sin\gamma + D_{\mathrm{initial}} - D_{\mathrm{react}} \\ Y_{\mathrm{cp3}} = -R_{\mathrm{turn}} \cos\gamma + R_{\mathrm{turn}} \end{cases} \quad [6]$$

The x and y coordinates of the 1st part of the safety zone edge ($X_{\mathrm{safe1}}$, $Y_{\mathrm{safe1}}$) are given by:

$$Y_{\mathrm{safe1}} = D_{\mathrm{width}} \quad [7]$$

The x and y coordinates of the 2nd part of the safety zone edge ($X_{\mathrm{safe2}}$, $Y_{\mathrm{safe2}}$) are given by:

$$(X_{\mathrm{safe2}} - D_{\mathrm{initial}} + D_{\mathrm{react}})^2 + (Y_{\mathrm{safe2}} - R_{\mathrm{turn}})^2 = {D_2}^2 + (D_{\mathrm{width}} - R_{\mathrm{turn}})^2 \quad [8]$$

The x and y coordinates of the 3rd part of the safety zone edge ($X_{\mathrm{safe3}}$, $Y_{\mathrm{safe3}}$) are given by:

$$(X_{\mathrm{safe3}} - D_{\mathrm{initial}} + D_{\mathrm{react}} - R_{\mathrm{device}} \cos\gamma + R_{\mathrm{turn}} \sin\gamma)^2 + (Y_{\mathrm{safe3}} - R_{\mathrm{turn}} + R_{\mathrm{device}} \sin\gamma + R_{\mathrm{turn}} \cos\gamma)^2 = {R_{\mathrm{device}}}^2 \quad [9]$$

For calculating the x and y coordinates of the 4th part of the safety zone edge ($X_{\mathrm{safe4}}$, $Y_{\mathrm{safe4}}$), the formula of involute curve from a circle of radius $R_{\mathrm{turn}}$ whose center is the origin (0, 0) with a tip of the string at ($R_{\mathrm{turn}}$, 0) is introduced as:

$$\begin{cases} x = R_{\mathrm{turn}} (\cos\theta + \theta \sin\theta) \\ y = R_{\mathrm{turn}} (\sin\theta - \theta \cos\theta) \end{cases}, 0 \le \theta \le \gamma \quad [10]$$

where $\theta$ denotes an angle for an unwrapped string from the circle in a counter-clockwise direction. By moving and rotating this point ($x$, $y$), we can calculate the 4th part of the safety zone edge ($X_{\mathrm{safe4}}$, $Y_{\mathrm{safe4}}$) as:

$$\begin{cases} X_{\mathrm{safe4}} = D_{\mathrm{initial}} - D_{\mathrm{react}} - x \sin\gamma + y \cos\gamma \\ Y_{\mathrm{safe4}} = R_{\mathrm{turn}} - x \cos\gamma + y \sin\gamma \end{cases} \quad [11]$$

From equations [1] to [11], the x and y coordinates of the crossing point of the escape path and the safety zone edge ($X_{\mathrm{cross}}$, $Y_{\mathrm{cross}}$) are given by a function of $D_{\mathrm{width}}$, $D_{\mathrm{attack}}$, $R_{\mathrm{device}}$, $D_{\mathrm{initial}}$, $D_{\mathrm{react}}$, $R_{\mathrm{turn}}$, and $\alpha+\beta$.

The prey can escape from the predator when the time required for the prey to enter the safety zone ($T_{prey}$) is shorter than the time required for the predator's capture device to reach that entry point ($T_{pred}$). Therefore, the prey is assumed to maximize the difference between the $T_{pred}$ and $T_{prey}$ ($T_{diff}$). To incorporate the time required for the prey to turn, $T_{prey}$ was divided into two phases: the fast-start phase, which includes the time for turning and acceleration ($T_1$), and the constant speed phase ($T_2$). This assumption is consistent with the previous studies [2-4] and was supported by our experiment (See Fig. S3). Therefore:

$$T_{prey} = T_1 + T_2 \quad [12]$$

For simplicity, the prey was assumed to end the fast-start phase at a certain displacement from the initial position in any $\alpha$ ($D_1$; the radius of the dotted circle in Fig. 1B) and to move at a constant speed $U_{prey}$ to cover the rest of the distance (toward the edge of the safety zone $\sqrt{X_{cross}{}^2 + Y_{cross}{}^2} - D_1$, plus the length of the body that is posterior to the center of mass $L_{prey}$). Because a larger $|\alpha|$ requires further turning prior to forward locomotion, which takes time [2, 5], and the initial velocity after turning was dependent on $|\alpha|$ in our experiment (See Fig. 3B), $T_1$ is given by a function of $|\alpha|$ [$T_1(|\alpha|)$]. Therefore, $T_{prey}$ can be expressed as:

$$T_{prey} = T_1(|\alpha|) + \frac{\sqrt{X_{cross}{}^2 + Y_{cross}{}^2} - D_1 + L_{prey}}{U_{prey}} \quad [13]$$

When the prey reaches the 1st part of the safety zone edge, $T_{pred}$ can be expressed as:

$$T_{pred} = \frac{D_{initial} + D_2 + D_{react} - X_{cross}}{U_{pred}} \quad [14]$$

When the prey reaches the 2nd part of the safety zone edge, $T_{pred}$ can be expressed as:

$$T_{pred} = \frac{D_{react} + R_{turn} \tan^{-1} \frac{D_2(Y_{cross} - R_{turn}) - (X_{cross} - D_{initial} + D_{react})(D_{width} - R_{turn})}{D_2(X_{cross} - D_{initial} + D_{react}) + (D_{width} - R_{turn})(Y_{cross} - R_{turn})}}{U_{pred}} \quad [15]$$

When the prey reaches the 3rd or 4th part of the safety zone edge, $T_{pred}$ can be expressed as:

$$T_{pred} = \frac{D_{initial} + D_{attack}}{U_{pred}} \quad [16]$$

From equations [1] to [16], we can calculate $T_{\mathrm{diff}}$ in response to the changes of $\alpha$ and $\beta$, from $D_1$, $D_{\mathrm{width}}$, $D_{\mathrm{attack}}$, $R_{\mathrm{device}}$, $D_{\mathrm{initial}}$, $D_{\mathrm{react}}$, $R_{\mathrm{turn}}$, $U_{\mathrm{prey}}$, $U_{\mathrm{pred}}$, and $T_1(|\alpha|)$. Given that the escape success is assumed to be dependent on $T_{\mathrm{diff}}$, the theoretically optimal ET can be expressed as:

$$\text{The optimal ET} = \underset{\alpha+\beta}{\operatorname{argmax}}(T_{\mathrm{diff}}) \qquad [17]$$

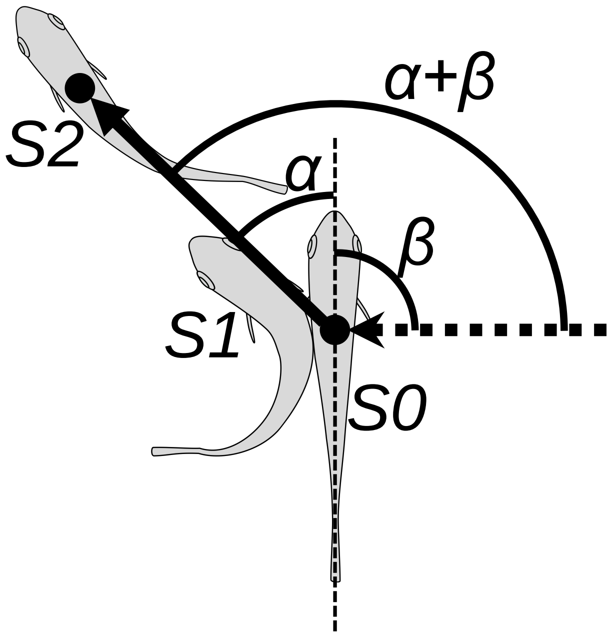


Fig. S1. Schematic drawing of angular variables. *Filled circle* position of the center of mass; *Dotted arrow* approach direction of the dummy predator; *S0* position of the fish at the onset of stage 1, *S1* position at the end of stage 1, *S2* position at the end of stage 2, $\alpha$ turn angle, $\beta$ initial orientation, $\alpha+\beta$ escape trajectory (ET).

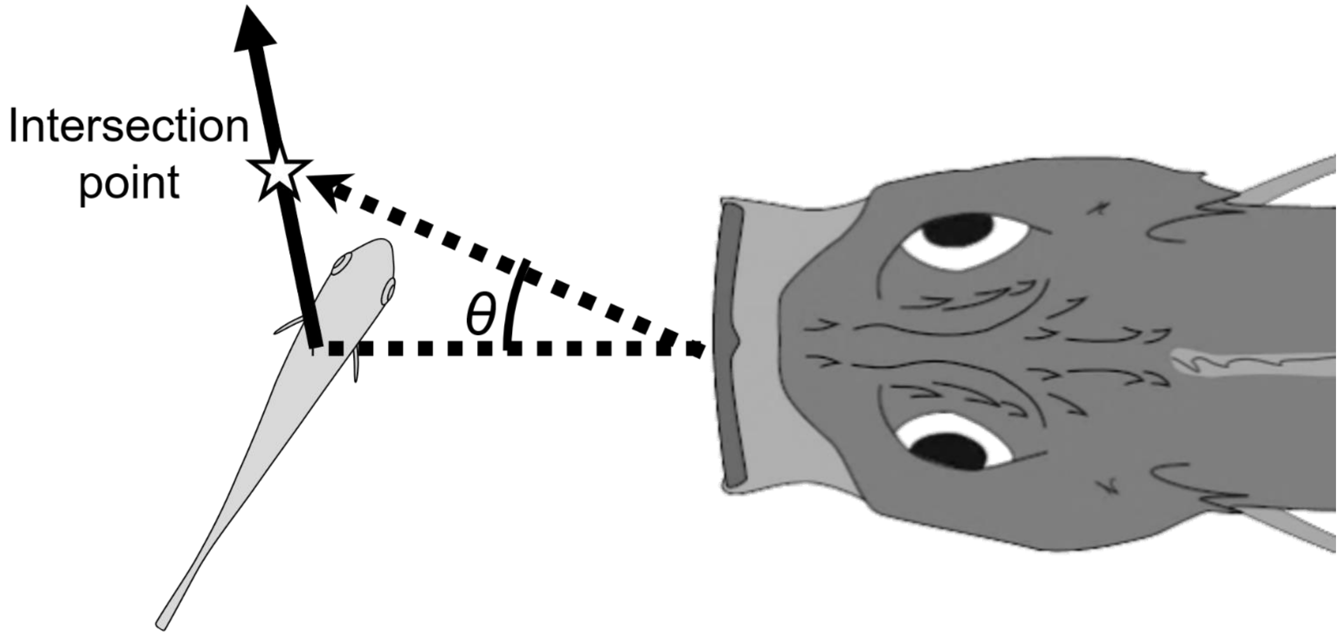


Fig. S2. Schematic drawing of how the adjusted angle of the predator ($\theta$) was measured. The intersection point is the crossing point between the trajectory of the prey's center of mass (CoM) and the trajectory of the predator's tip of the mouth. The adjusted angle is defined as the angle between the line passing through the predator's tip of the mouth and the prey's CoM at the onset of the prey's escape response, and the line passing through the predator's tip of the mouth at the onset of the escape response and the intersection point.

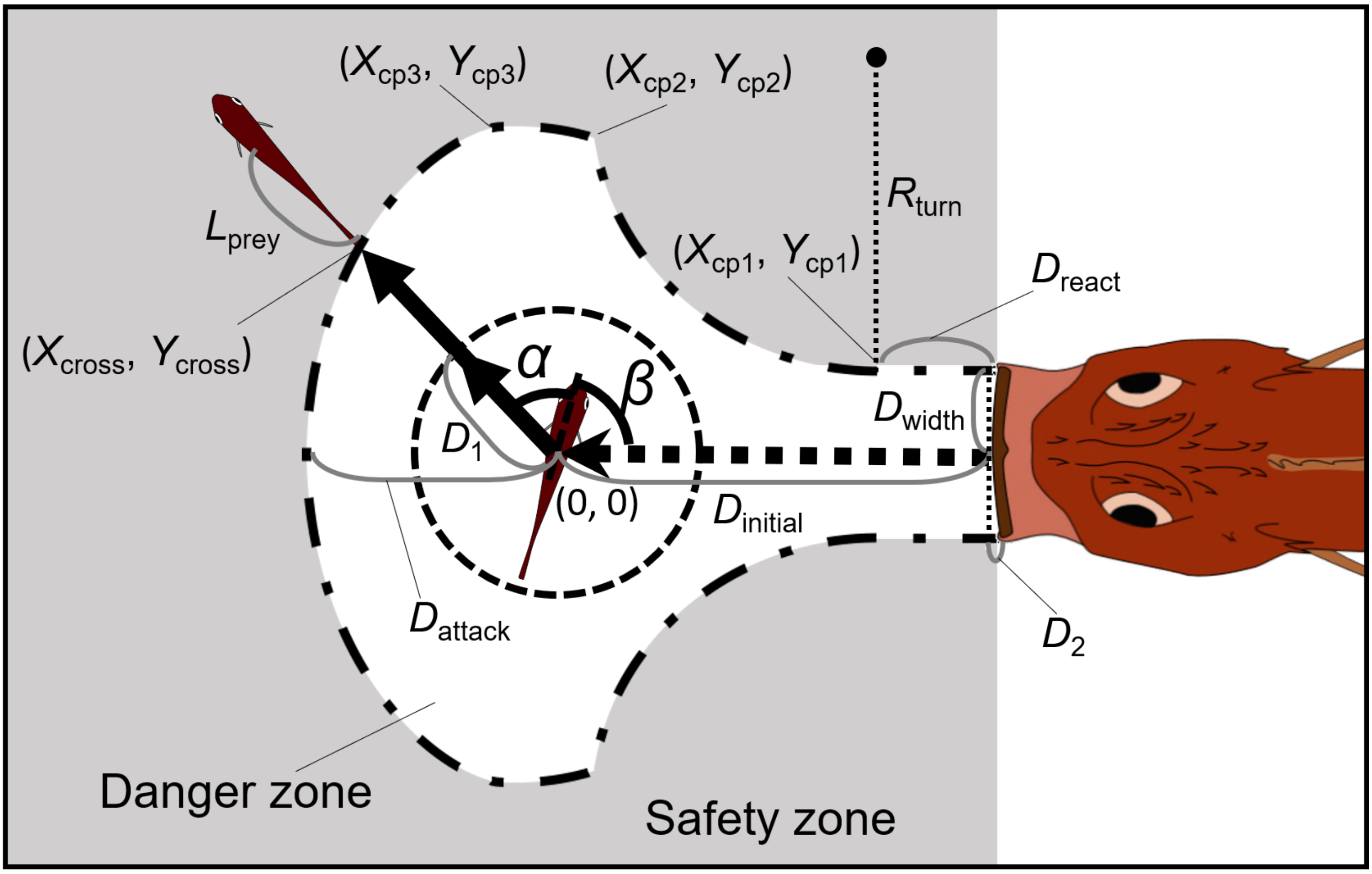


Fig. S3. The geometric model modified from Corcoran & Conner (2016) [1]. Two factors are added to Corcoran's model: the endpoint of the predator attack, and the time required for the prey to turn. See SI text for details of the definitions of the variables and mathematical formulas.

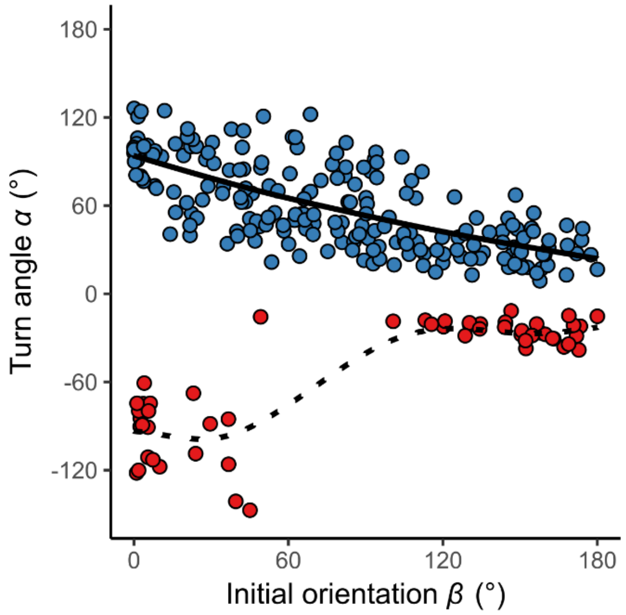


Fig. S4. Relationship between initial orientation $\beta$ and turn angle $\alpha$ in the experiment. Different colors represent the away (blue) and toward (red) responses. Solid and dotted lines are estimated by the generalized additive mixed model (GAMM) [n=264 (208 away and 56 toward responses) from 23 individuals].

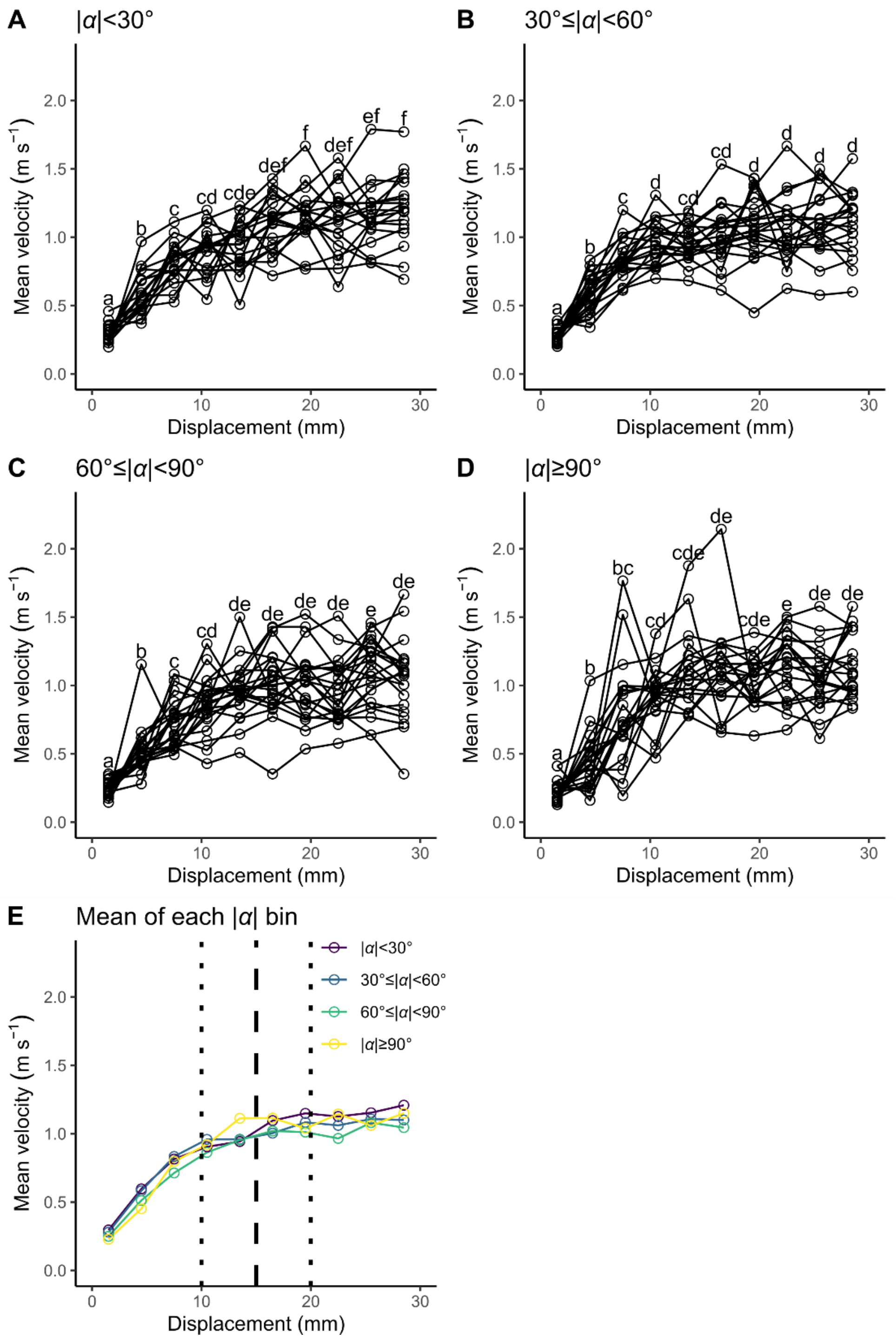


Fig. S5. Relationship between displacement from the initial position (3-mm intervals: 0–3, 3–6, ..., and 27–30 mm) and mean velocity during the displacement for each turn angle (|α|) bin. Unfilled circles denote the mean value for each individual. Different lowercase letters represent significant differences according to the paired *t*-test with Bonferroni's correction ($P$<0.05). (A) |α|<30°. (B) 30°≤|α|<60°. (C) 60°≤|α|<90°. (D) |α|≥90°. (E) Mean of the individual mean value for each |α| bin. Vertical dashed line represents the cut-off distance of 15 mm used in this study, and vertical dotted lines represent the other cut-off distances tested in this study (Tables S2 and S3) (n=23 individuals).

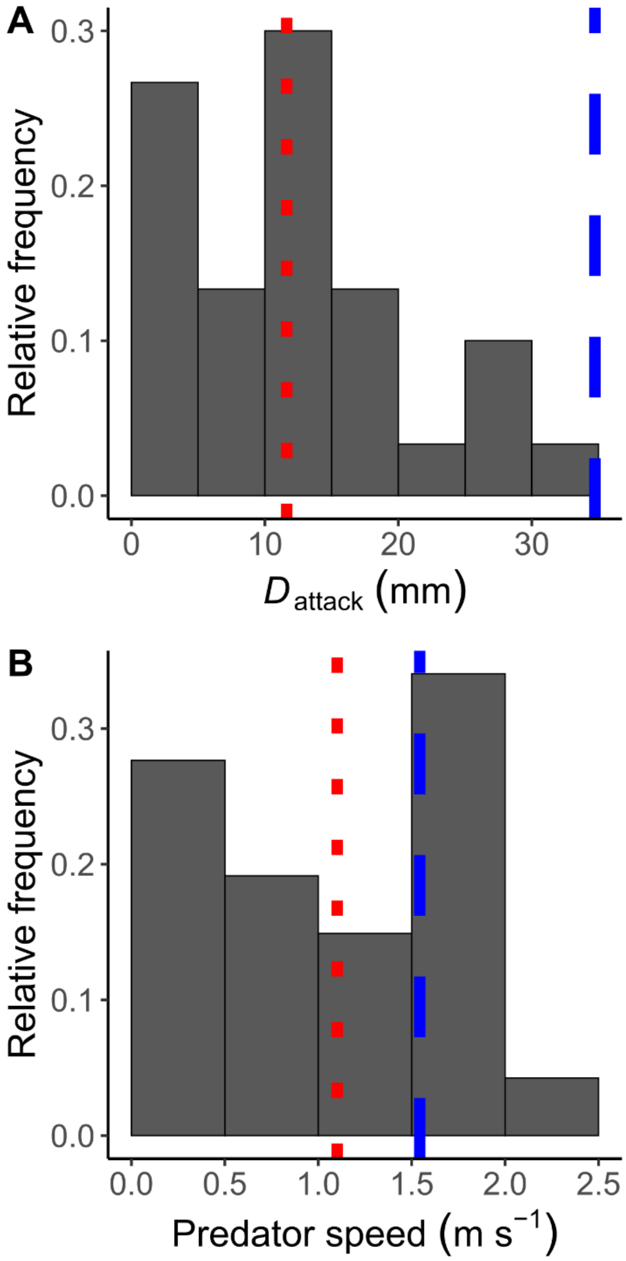


Fig. S6. Predator *Sebastiscus marmoratus* attack parameters. (A) Histogram of the distance between the prey's initial position and the predator's mouth position at the onset of the mouth closing ($D_{\mathrm{attack}}$) (n=30 from 7 individuals). (B) Histogram of the speed of the real predator (n=47 from 7 individuals). Both figures are based on reanalysis of data from Kimura and Kawabata [6]. Vertical dashed blue lines represent the optimal values independently estimated in this study, and vertical dotted red lines represent the mean values of the real predator.

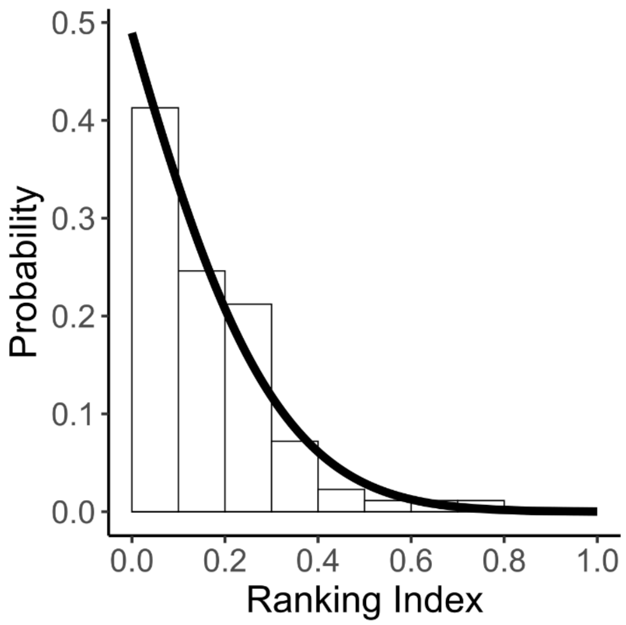


Fig. S7. Histogram of the ranking index, where 0 indicates that the real fish chose the theoretically optimal escape trajectory (ET) and 1 indicates that the real fish chose the theoretically worst ET. The solid line is the density probability function of the truncated normal distribution (n=264 from 23 individuals).

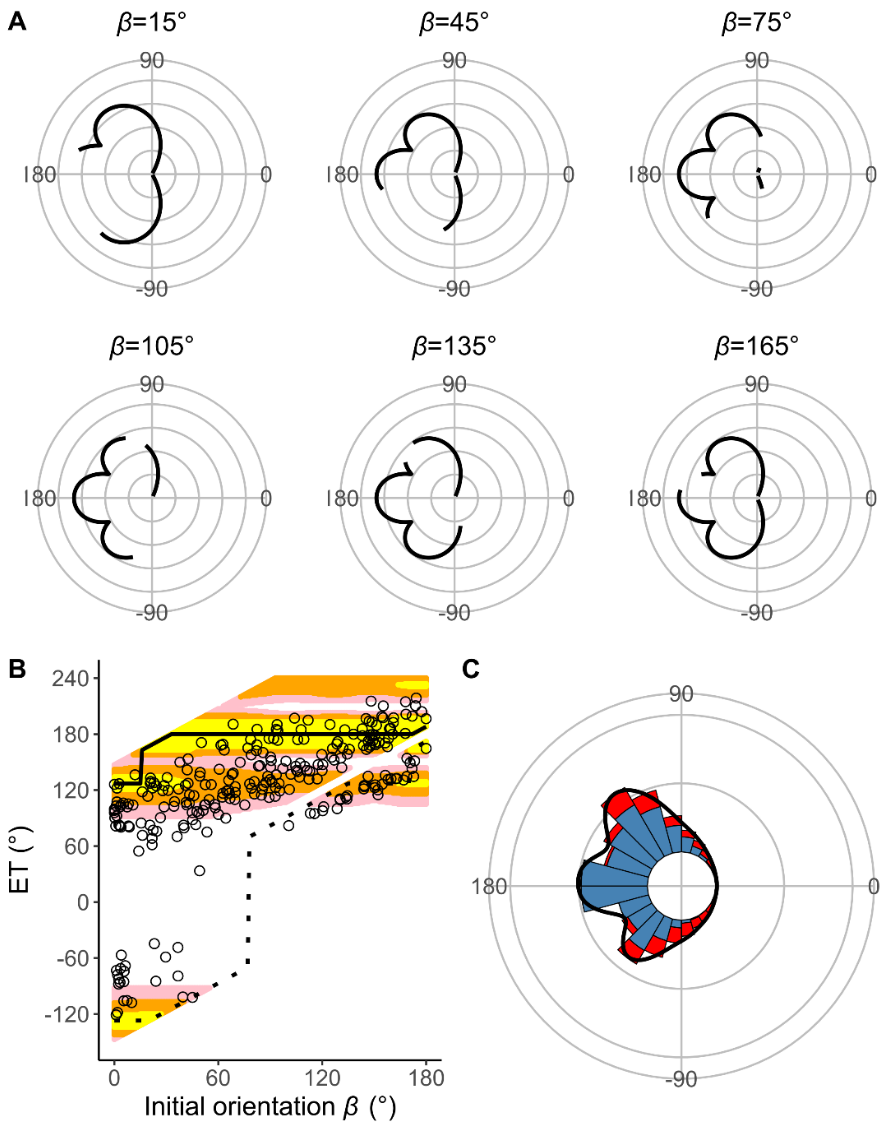


Fig. S8. Estimates of the model with $D_{attack}$ (the distance between the prey's initial position and the endpoint of the predator attack) and without $T_1(|\alpha|)$ (the relationship between the absolute value of the turn angle $|\alpha|$ and the time required for a 15-mm displacement from the initial position, or the time required for prey to turn). (A) Circular plots of the time difference between the prey and predator $T_{diff}$ in different initial orientations $\beta$. The time difference of the best escape trajectory (ET) was regarded as 10 ms, and the relative time differences between 0 and 10 ms are shown by solid lines. Areas without solid lines indicate that either the time difference is below 0 or the fish cannot go to that ET because of the constraint on the possible range of $|\alpha|$. Concentric circles represent 3 ms. (B) Relationship between the initial orientation $\beta$ and ET. Solid and dotted lines represent the best-estimated away and toward responses, respectively. Different colors represent the top 10%, 25%, and 40% quantiles of the time difference between the prey and predator within all possible ETs. (C) Circular histogram of the theoretical ETs, estimated by a Monte Carlo simulation. The probability of selection of an ET was determined by the truncated normal distribution of the optimal ranking index. This process was repeated 1000 times to estimate the frequency distribution of the theoretical ETs. Colors in the bars represent the away (blue) or toward (red) responses. Black lines represent the kernel probability density function. Concentric circles represent 10 % of the total sample sizes, the bin intervals are 15°, and the bandwidths of the kernel are 50. The predator's approach direction is represented by 0° (n=264 from 23 individuals for experimental data, and n=264000 for Monte Carlo simulation).

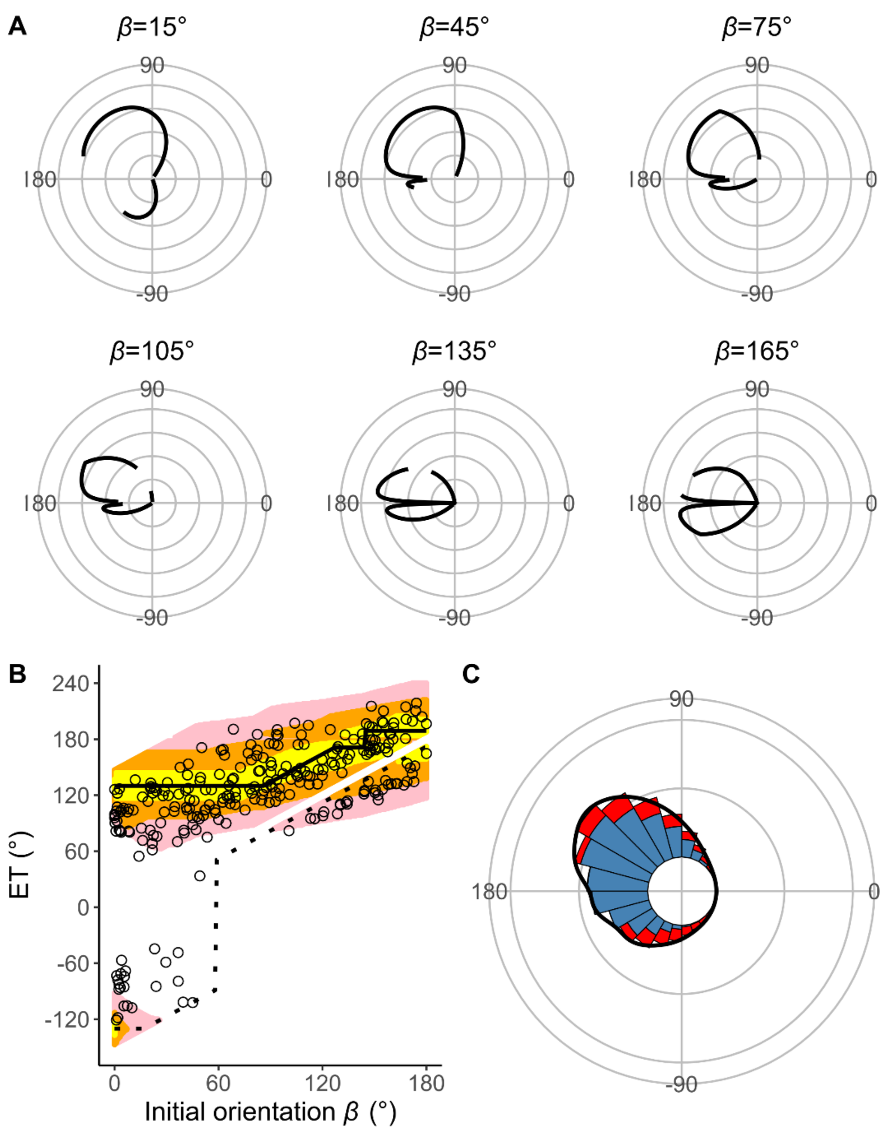


Fig. S9. Estimates of the model with $T_1(|\alpha|)$ (the relationship between the absolute value of the turn angle $|\alpha|$ and the time required for a 15-mm displacement from the initial position, or the time required for the prey to turn) and without $D_{\mathrm{attack}}$ (the distance between the prey's initial position and the endpoint of the predator attack). (A) Circular plots of the time difference between the prey and predator $T_{\mathrm{diff}}$ in different initial orientations $\beta$. The time difference of the best escape trajectory (ET) was regarded as 10 ms, and the relative time differences between 0 and 10 ms are shown by solid lines. Areas without solid lines indicate that either the time difference is below 0 or the fish cannot go to that ET because of the constraint on the possible range of $|\alpha|$. Concentric circles represent 3 ms. (B) Relationship between the initial orientation $\beta$ and ET. Solid and dotted lines represent the best-estimated away and toward responses, respectively. Different colors represent the top 10%, 25%, and 40% quantiles of the time difference between the prey and predator within all possible ETs. (C) Circular histogram of the theoretical ETs, estimated by a Monte Carlo simulation. The probability of selection of an ET was determined by the truncated normal distribution of the optimal ranking index. This process was repeated 1000 times to estimate the frequency distribution of the theoretical ETs. Colors in the bars represent the away (blue) or toward (red) responses. Black lines represent the kernel probability density function. Concentric circles represent 10 % of the total sample sizes, the bin intervals are 15°, and the bandwidths of the kernel are 50. The predator's approach direction is represented by 0° (n=264 from 23 individuals for experimental data, and n=264000 for Monte Carlo simulation).

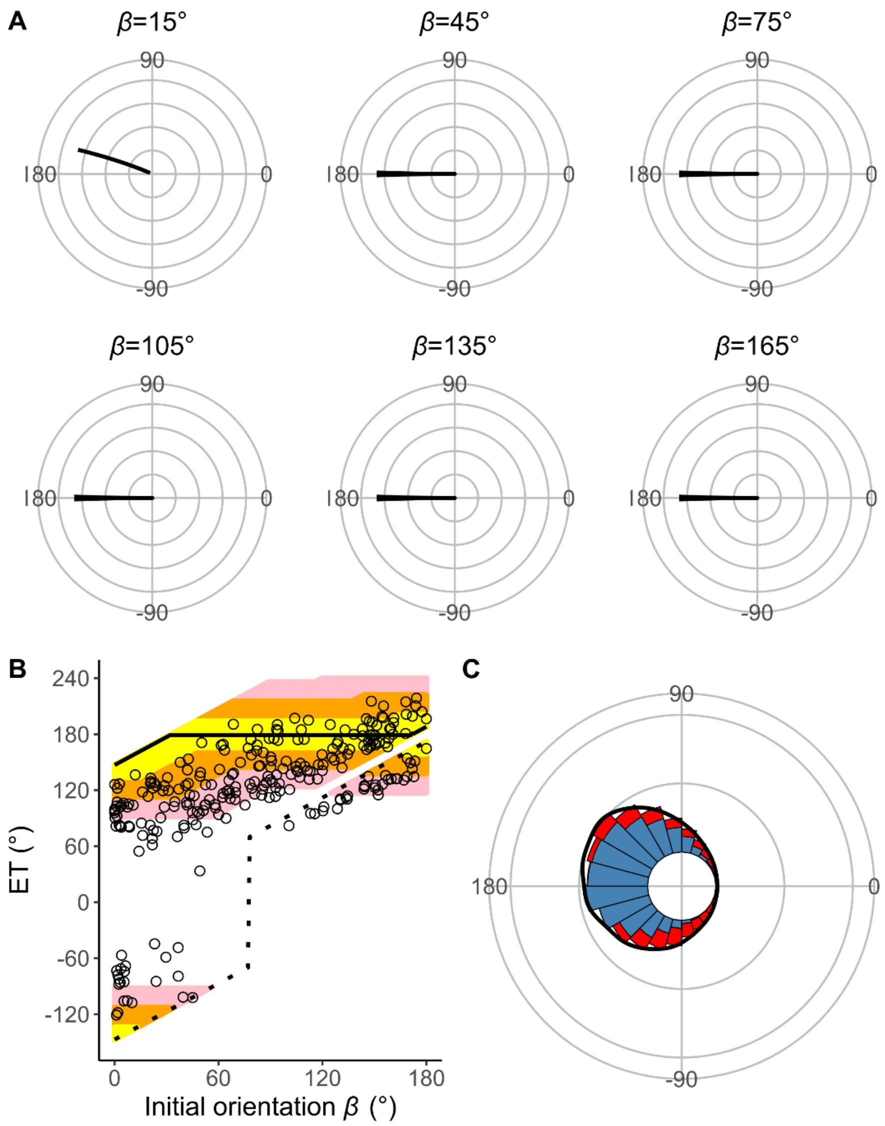


Fig. S10. Estimates of the model that includes neither $D_{\text{attack}}$ (the distance between the prey's initial position and the endpoint of the predator attack) nor $T_1(|\alpha|)$ (the relationship between the absolute value of the turn angle $|\alpha|$ and the time required for a 15-mm displacement from the initial position, or the time required for the prey to turn). (A) Circular plots of the time difference between the prey and predator $T_{\text{diff}}$ in different initial orientations $\beta$. The time difference of the best escape trajectory (ET) was regarded as 10 ms, and the relative time differences between 0 and 10 ms are shown by solid lines. Areas without solid lines indicate that either the time difference is below 0 or the fish cannot go to that ET because of the constraint on the possible range of $|\alpha|$. Concentric circles represent 3 ms. (B) Relationship between the initial orientation $\beta$ and ET. Solid and dotted lines represent the best-estimated away and toward responses, respectively. Different colors represent the top 10%, 25%, and 40% quantiles of the time difference between the prey and predator within all possible ETs. (C) Circular histogram of the theoretical ETs, estimated by a Monte Carlo simulation. The probability of selection of an ET was determined by the truncated normal distribution of the optimal ranking index. This process was repeated 1000 times to estimate the frequency distribution of the theoretical ETs. Colors in the bars represent the away (blue) or toward (red) responses. Black lines represent the kernel probability density function. Concentric circles represent 10 % of the total sample sizes, the bin intervals are 15°, and the bandwidths of the kernel are 50. The predator's approach direction is represented by 0° (n=264 from 23 individuals for experimental data, and n=264000 for Monte Carlo simulation).

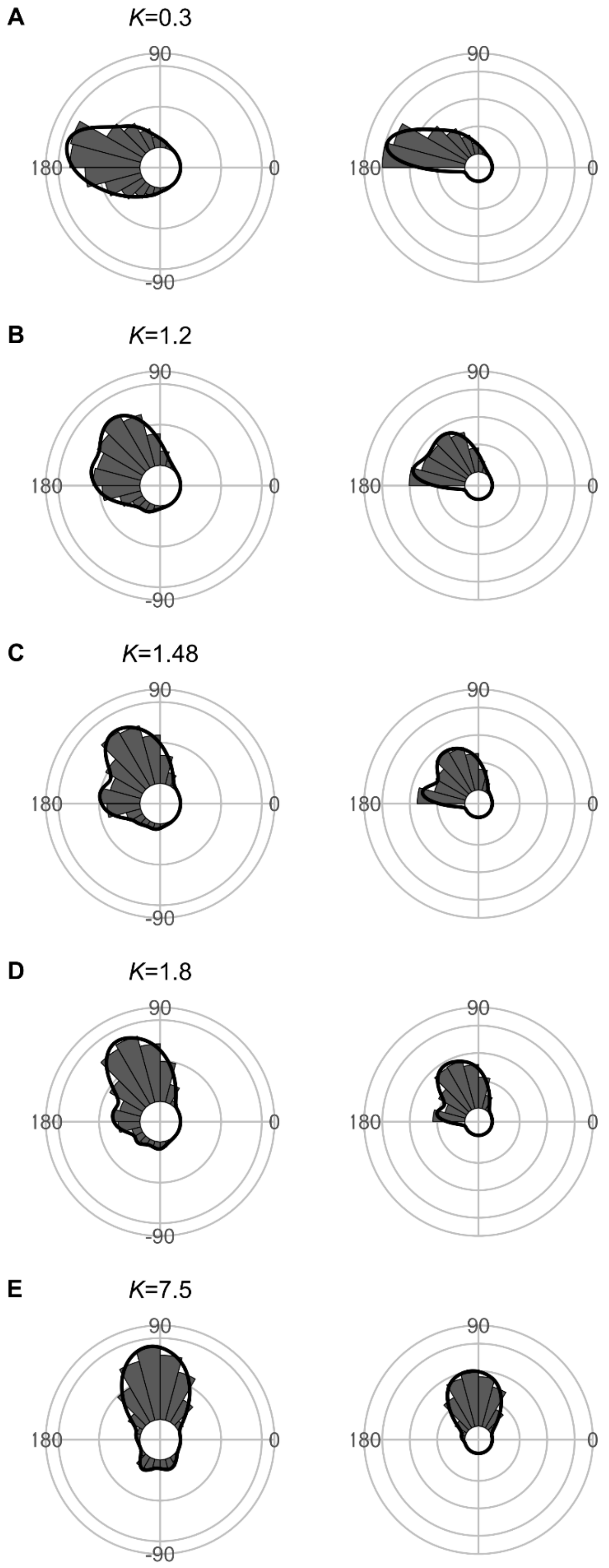


Fig. S11. Effect of predator speed $U_{\text{pred}}$ ($K=U_{\text{pred}}/U_{\text{prey}}$) on the theoretical distribution of escape trajectories (ET, left panel; $\text{ET}_{\text{semi}}$, right panel). Circular histograms of the theoretical escape trajectories were estimated by a Monte Carlo simulation of the geometric model. $\text{ET}_{\text{semi}}$ denotes the angle for escape trajectory ranging from 0° (directly toward the threat) to 180° (opposite to the threat), thereby using only one semicircle. The other parameter values are the same as the values used for explaining the escape response of *Pagrus major*.

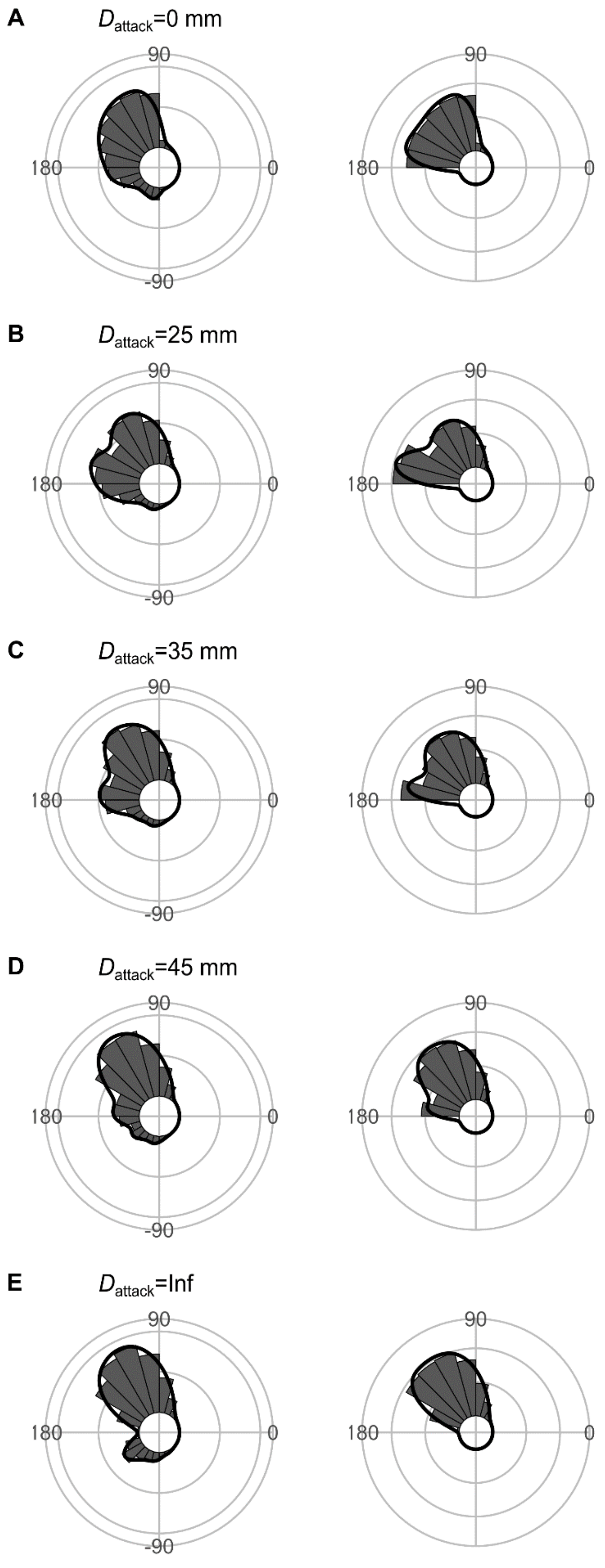


Fig. S12. Effect of $D_{attack}$ (the distance between the prey's initial position and the endpoint of the predator attack) on the theoretical distribution of escape trajectories (ET, left panel; $ET_{semi}$, right panel). Circular histograms of the theoretical escape trajectories were estimated by a Monte Carlo simulation of the geometric model. The other parameter values are the same as the values used for explaining the escape response of *Pagrus major*.

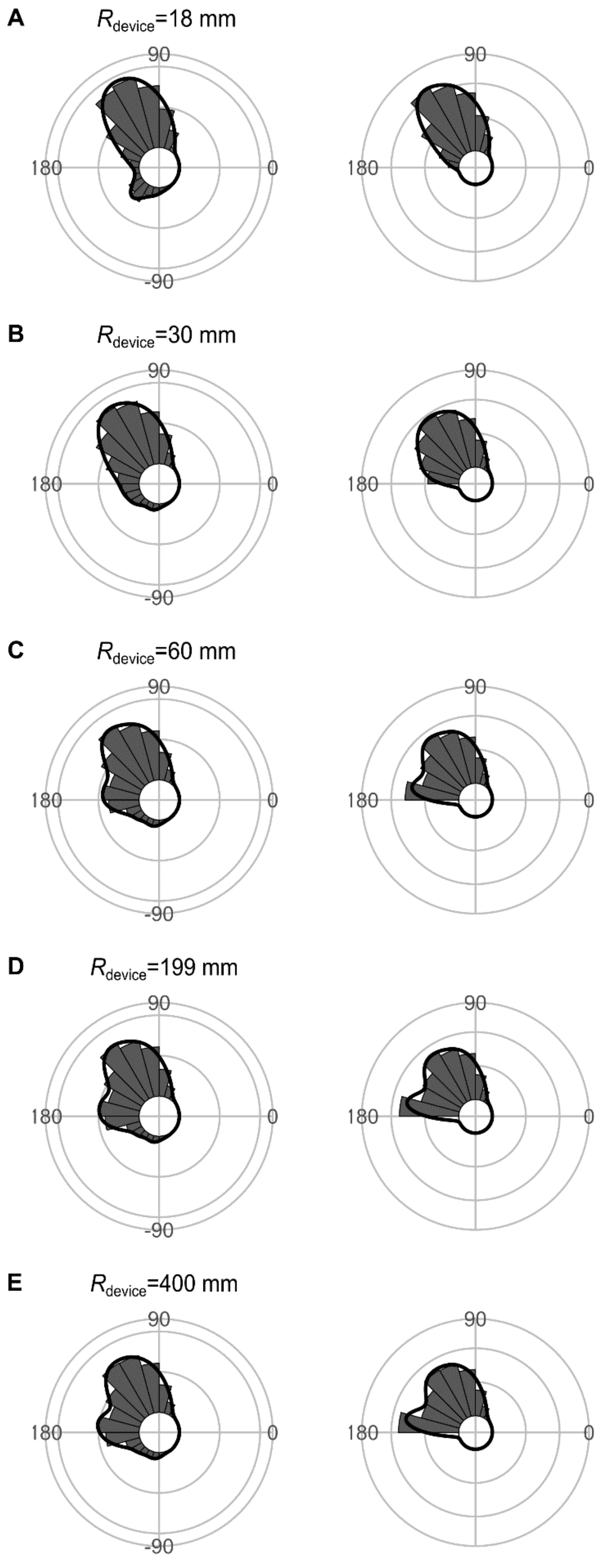


Fig. S13. Effect of $R_{device}$ (the radius for the shape of the predator's capture device at the moment of attack, which is approximated as an arc) on the theoretical distribution of escape trajectories (ET, left panel; $ET_{semi}$, right panel). Circular histograms of the theoretical escape trajectories were estimated by a Monte Carlo simulation of the geometric model. The other parameter values are the same as the values used for explaining the escape response of *Pagrus major*.

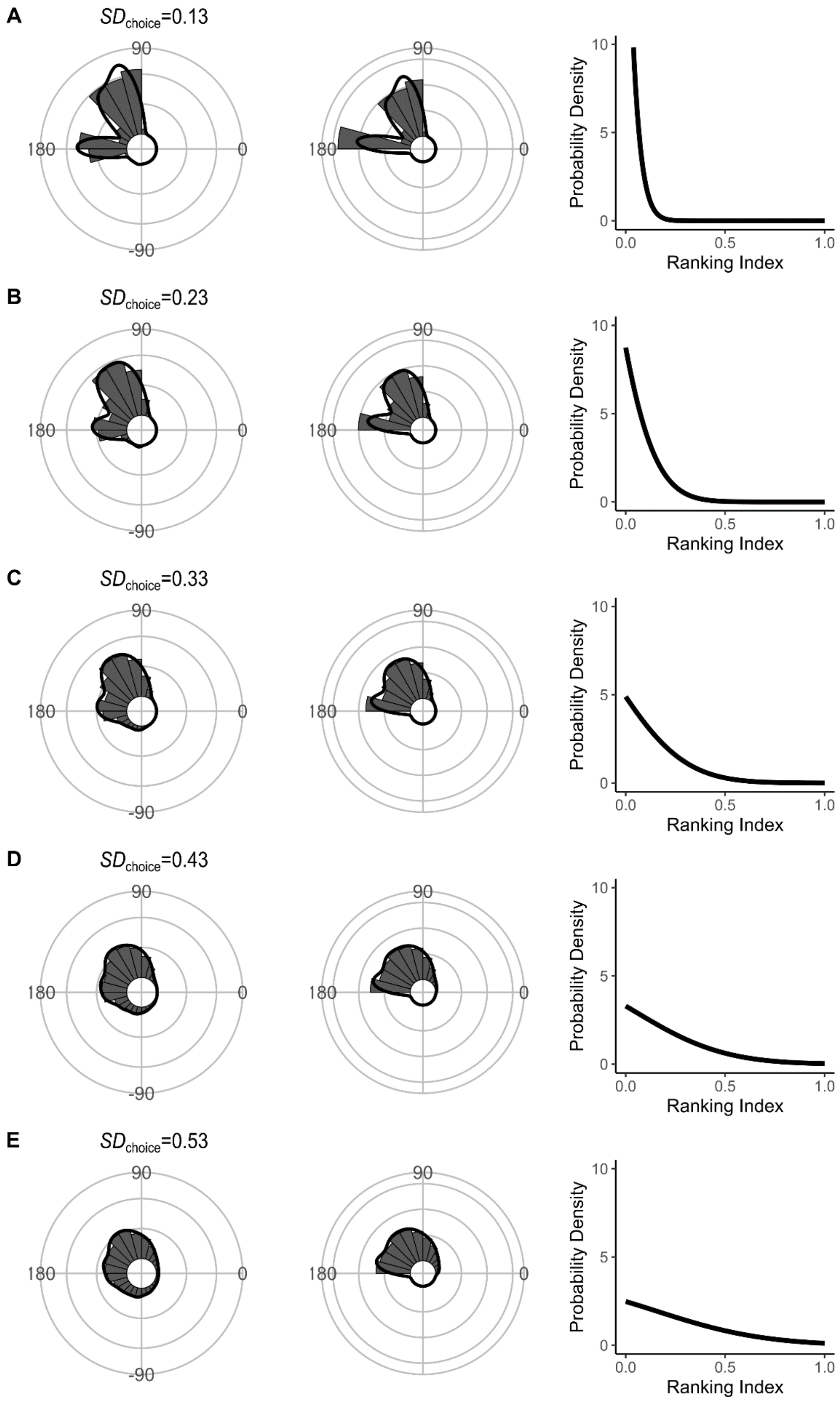


Fig. S14. Effect of $SD_{choice}$ [s.d. of the truncated normal distribution for ET choice from the continuum of the optimal ET (ranking index=0) and worst ET (ranking index=1)] on the theoretical distribution of escape trajectories (ET, left panel; $ET_{semi}$, middle panel). Circular histograms of the theoretical escape trajectories were estimated by a Monte Carlo simulation of the geometric model. The other parameter values are the same as the values used for explaining the escape response of *Pagrus major*.

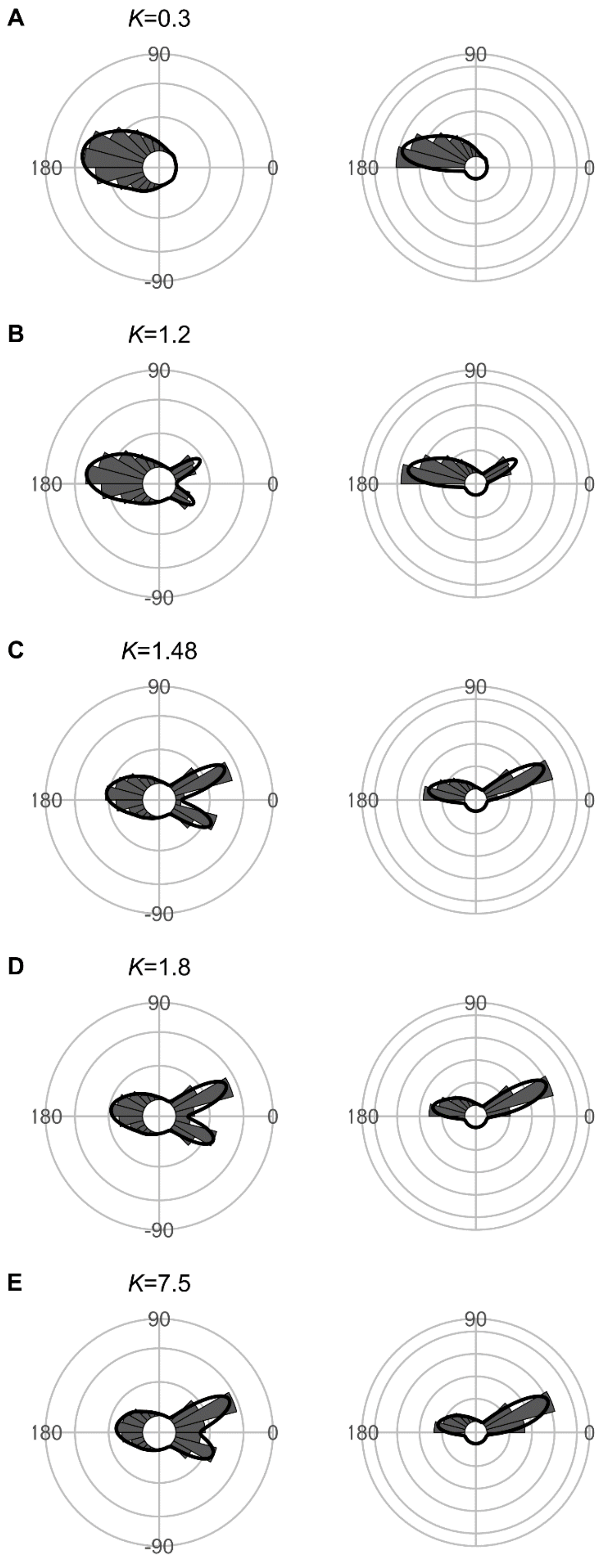


Fig. S15. Effect of predator speed $U_{pred}$ ($K=U_{pred}/U_{prey}$) on the theoretical distribution of escape trajectories (ET, left panel; $ET_{semi}$, right panel). Circular histograms of the theoretical escape trajectories were estimated by a Monte Carlo simulation of the geometric model where the predator can adjust its approach path. $D_{initial}$ is 130 mm, $D_{react}$ is 70 mm, $R_{turn}$ is 12 mm, $D_{attack}$ is 400 mm, and the other parameter values are the same as the values used for explaining the escape response of *Pagrus major*.

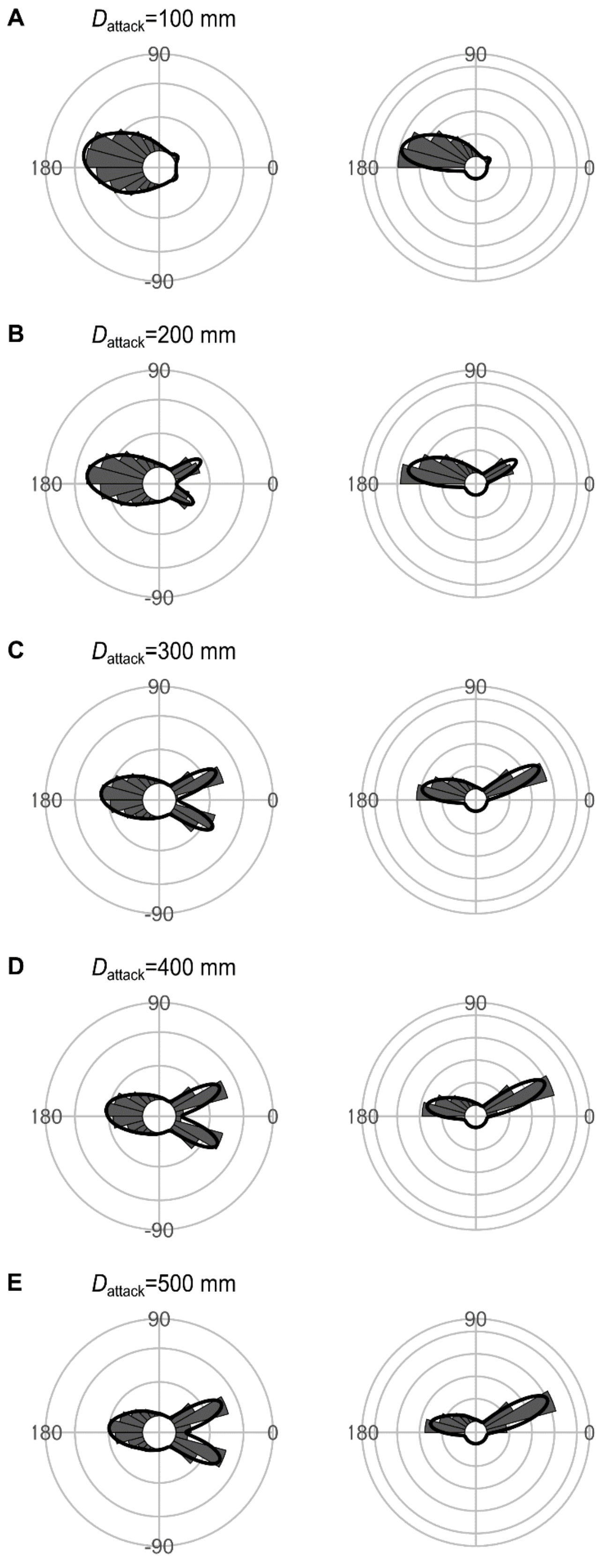


Fig. S16. Effect of $D_{attack}$ (the distance between the prey's initial position and the endpoint of the predator attack) on the theoretical distribution of escape trajectories (ET, left panel; $ET_{semi}$, right panel). Circular histograms of the theoretical escape trajectories were estimated by a Monte Carlo simulation of the geometric model where the predator can adjust its approach path. $D_{initial}$ is 130 mm, $D_{react}$ is 70 mm, $R_{turn}$ is 12 mm, and the other parameter values are the same as the values used for explaining the escape response of *Pagrus major*.

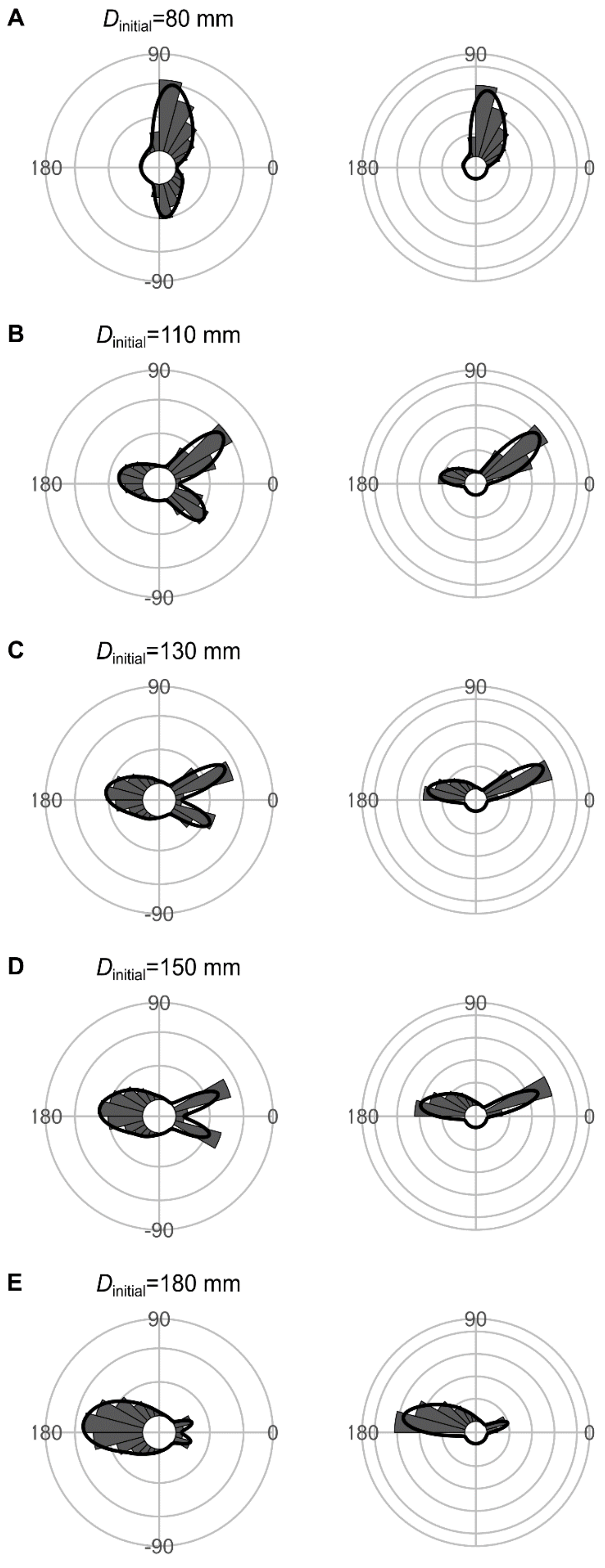


Fig. S17. Effect of $D_{initial}$ (the distance between the prey and the predator at the onset of the prey's escape response) on the theoretical distribution of escape trajectories (ET, left panel; $ET_{semi}$, right panel). Circular histograms of the theoretical escape trajectories were estimated by a Monte Carlo simulation of the geometric model where the predator can adjust its approach path. $D_{react}$ is 70 mm, $R_{turn}$ is 12 mm, $D_{attack}$ is 400 mm, and the other parameter values are the same as the values used for explaining the escape response of *Pagrus major*.

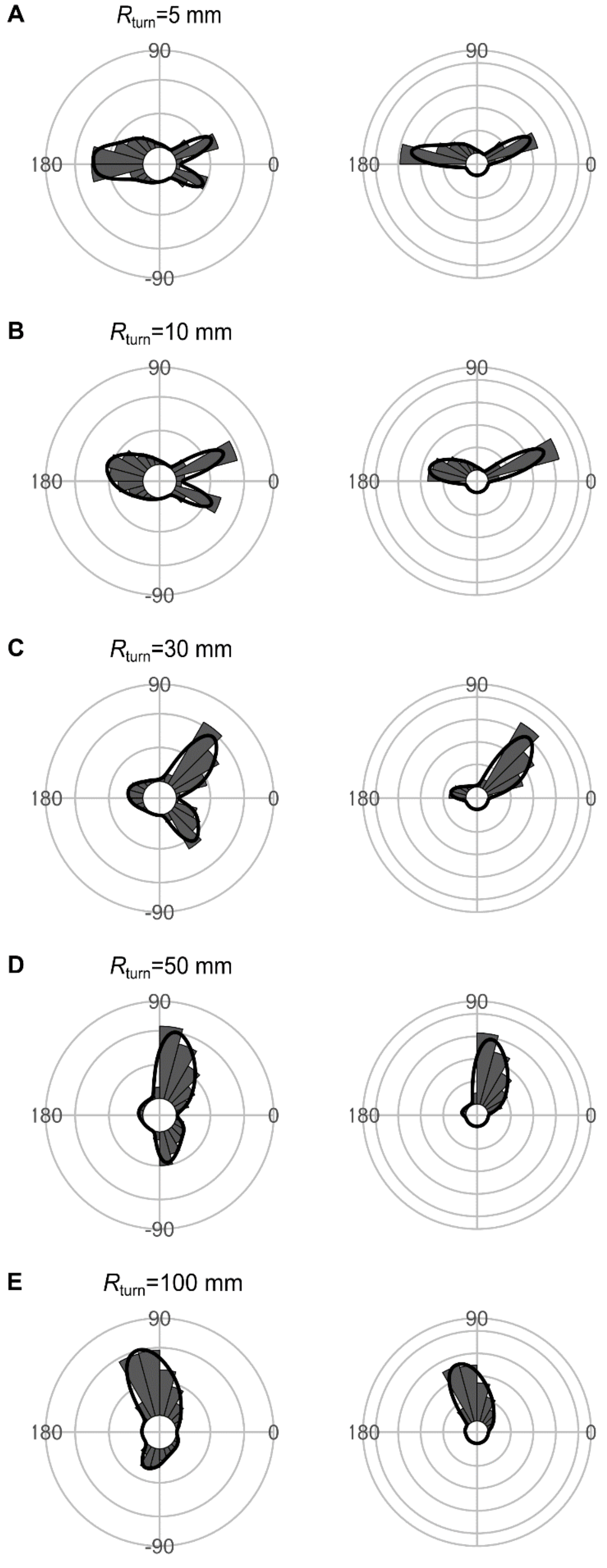


Fig. S18. Effect of the minimum turning radius of the predator $R_{turn}$ on the theoretical distribution of escape trajectories (ET, left panel; $ET_{semi}$, right panel). Circular histograms of the theoretical escape trajectories were estimated by a Monte Carlo simulation of the geometric model where the predator can adjust its approach path. $D_{initial}$ is 130 mm, $D_{react}$ is 70 mm, $D_{attack}$ is 400 mm, and the other parameter values are the same as the values used for explaining the escape response of *Pagrus major*.

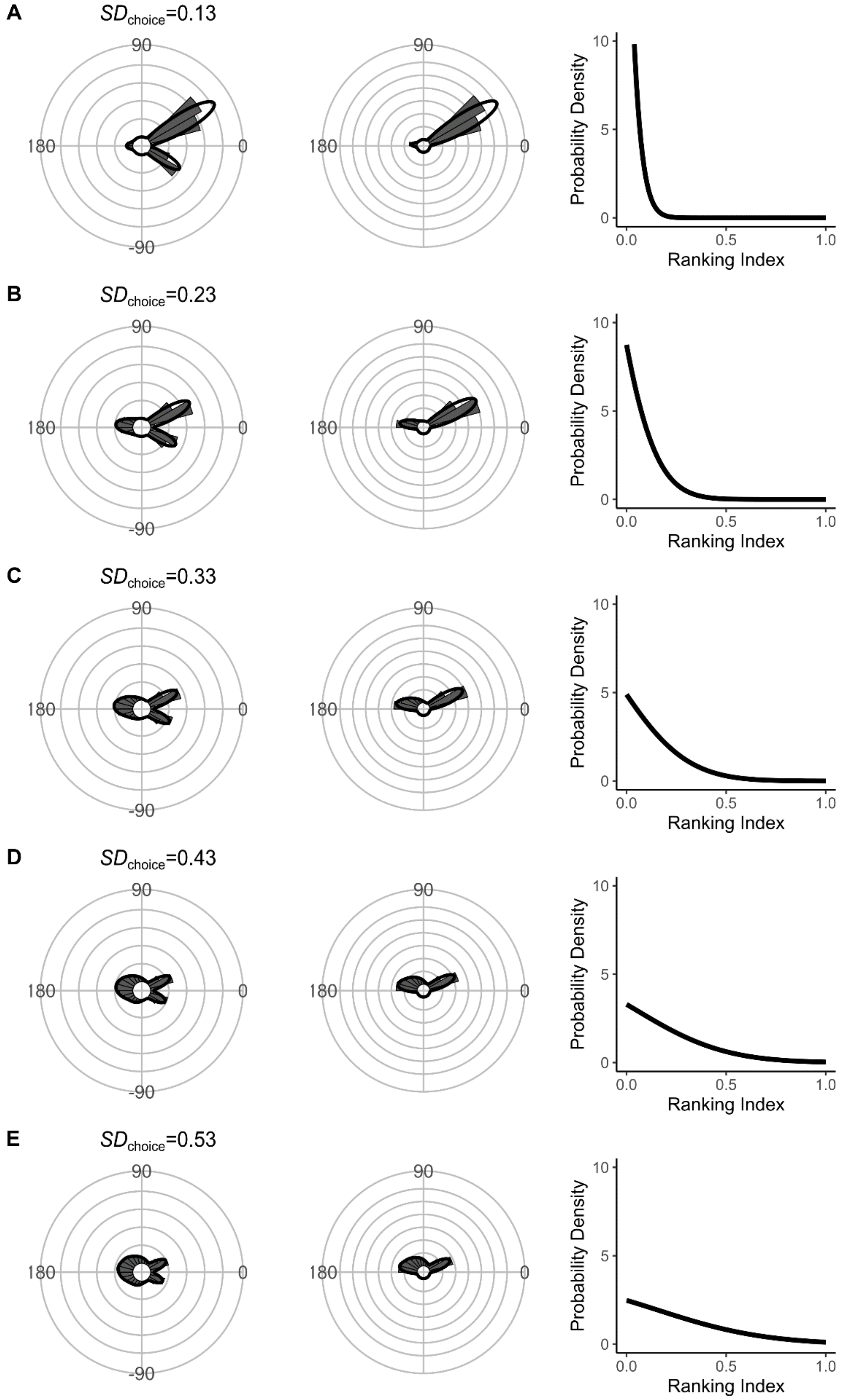


Fig. S19. Effect of $SD_{\mathrm{choice}}$ [s.d. of the truncated normal distribution for ET choice from the continuum of the optimal ET (ranking index=0) and worst ET (ranking index=1)] on the theoretical distribution of escape trajectories (ET, left panel; $\mathrm{ET}_{\mathrm{semi}}$, middle panel). Circular histograms of the theoretical escape trajectories were estimated by a Monte Carlo simulation of the geometric model where the predator can adjust its approach path. $D_{\mathrm{initial}}$ is 130 mm, $D_{\mathrm{react}}$ is 70 mm, $D_{\mathrm{attack}}$ is 400 mm, and the other parameter values are the same as the values used for explaining the escape response of *Pagrus major*.

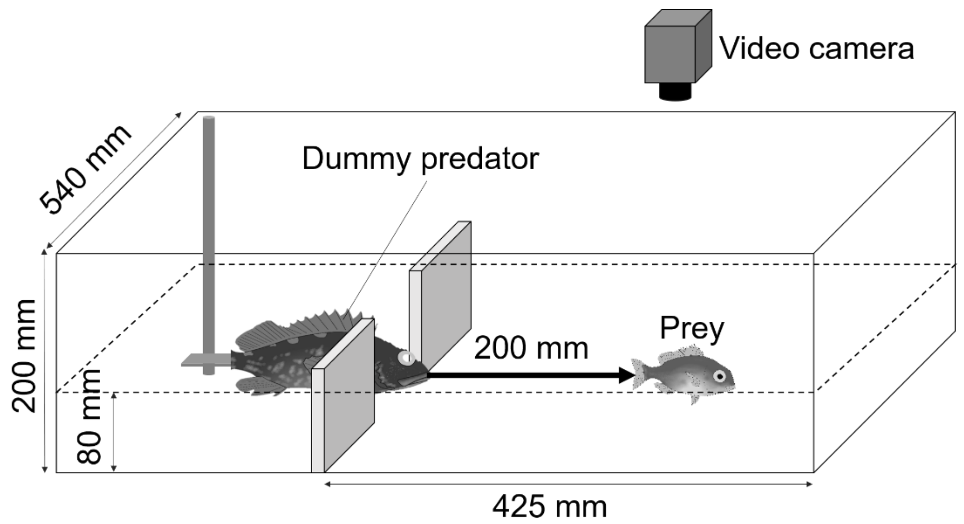


Fig. S20. Sketch of the experimental apparatus for measuring the escape response of prey fish *Pagrus major*.

Table S1. Akaike information criterion (AIC) for 1–9 Gaussian mixture models to estimate the empirical ET distribution (n=264 from 23 individuals)

| Number of peaks | AIC | ΔAIC |
|---|---|---|
| **3** | 2777.1 | 0.0 |
| 4 | 2781.0 | 3.9 |
| 2 | 2784.1 | 6.9 |
| 5 | 2787.1 | 10.0 |
| 6 | 2791.1 | 14.0 |
| 7 | 2797.1 | 20.0 |
| 8 | 2798.9 | 21.8 |
| 9 | 2799.2 | 22.1 |
| 1 | 2855.7 | 78.6 |

The best model is shown in bold.

Table S2. Widely applicable or Watanabe–Akaike information criterion (WAIC) for each model to estimate the relationship between the absolute value of the turn angle and the time required for a displacement of 10 or 20 mm from the initial position (n=264 from 23 individuals)

| Length of displacement | WAIC | ΔWAIC |
|---|---|---|
| 10 mm | | |
| **Piecewise linear** | 1239.7 | 0 |
| Linear | 1259.0 | 19.3 |
| Constant | 1524.4 | 284.7 |
| 20 mm | | |
| **Piecewise linear** | 1543.3 | 0 |
| Linear | 1547.0 | 3.7 |
| Constant | 1689.7 | 146.4 |

The best models are shown in bold.

Table S3. Comparison of the distribution of escape trajectories (ETs) between the model prediction (n=264 per simulation ×1000 times ×2 distances) and experimental data (n=264) using the two-sample Kuiper test

| Distance for the fast-start phase | Median Kuiper's $V$ | Median $P$ | Rate of $P > 0.05$ |
|---|---|---|---|
| 10 mm | | | |
| With both $D_{attack}$ and $T_1(\|\alpha\|)$ | 0.10 | 0.63 | 0.99 |
| With $D_{attack}$ and without $T_1(\|\alpha\|)$ | 0.25 | < 0.01 | 0.00 |
| Without $D_{attack}$ and with $T_1(\|\alpha\|)$ | 0.17 | < 0.05 | 0.25 |
| Neither $D_{attack}$ nor $T_1(\|\alpha\|)$ | 0.28 | < 0.01 | 0.00 |
| 20 mm | | | |
| With both $D_{attack}$ and $T_1(\|\alpha\|)$ | 0.11 | 0.44 | 0.96 |
| With $D_{attack}$ and without $T_1(\|\alpha\|)$ | 0.25 | < 0.01 | 0.00 |
| Without $D_{attack}$ and with $T_1(\|\alpha\|)$ | 0.15 | < 0.05 | 0.46 |
| Neither $D_{attack}$ nor $T_1(\|\alpha\|)$ | 0.28 | < 0.01 | 0.00 |

The distance for the fast-start phase was regarded as either 10 or 20 mm. $D_{attack}$, distance between the prey's initial position and the endpoint of the predator attack; $T_1(|\alpha|)$, relationship between the absolute value of the turn angle and the time required for a 15-mm displacement from the initial position (i.e., the time required for the prey to turn).

### SI References


1. Corcoran AJ, Conner WE. How moths escape bats: predicting outcomes of predator-prey interactions. J Exp Biol. 2016;219:2704-15. doi: 10.1242/jeb.137638.
2. Domenici P, Blake RW. The kinematics and performance of the escape response in the angelfish (*Pterophyllum eimekei*). J Exp Biol. 1991;156:187-205.
3. Danos N, Lauder GV. Challenging zebrafish escape responses by increasing water viscosity. J Exp Biol. 2012;215(Pt 11):1854-62. doi: 10.1242/jeb.068957.
4. Fleuren M, van Leeuwen JL, Quicazan-Rubio EM, Pieters RPM, Pollux BJA, Voesenek CJ. Three-dimensional analysis of the fast-start escape response of the least killifish, *Heterandria formosa*. J Exp Biol. 2018;221(Pt 7). doi: 10.1242/jeb.168609.
5. Ellerby DJ, Altringham JD. Spatial variation in fast muscle function of the rainbow trout *Oncorhynchus mykiss* during fast-starts and sprinting. J Exp Biol. 2001;204(13):2239-50.
6. Kimura H, Kawabata Y. Effect of initial body orientation on escape probability of prey fish escaping from predators. Biol Open. 2018;7(7):bio023812. doi: 10.1242/bio.023812.
